# Flexibility of intrinsically disordered degrons in AUX/IAA proteins reinforces auxin co-receptor assemblies

**DOI:** 10.1101/787770

**Authors:** Michael Niemeyer, Elena Moreno Castillo, Christian H. Ihling, Claudio Iacobucci, Verona Wilde, Antje Hellmuth, Wolfgang Hoehenwarter, Sophia L. Samodelov, Matias D. Zurbriggen, Panagiotis L. Kastritis, Andrea Sinz, Luz Irina A. Calderón Villalobos

**Affiliations:** Department of Molecular Signal Processing, Leibniz Institute of Plant Biochemistry (IPB), Weinberg 3, D-06120 Halle (Saale), Germany; Department of Pharmaceutical Chemistry & Bioanalytics, Institute of Pharmacy, Martin-Luther University Halle-Wittenberg, Charles Tanford Protein Center, Kurt-Mothes-Str. 3a, D-06120 Halle (Saale), Germany; Proteome Analytics Research Group Leibniz Institute of Plant Biochemistry (IPB), Weinberg 3, D-06120 Halle (Saale), Germany; Institute of Synthetic Biology & Cluster of Excellence on Plant Science (CEPLAS), Heinrich-Heine University of Düsseldorf, Universitätsstrasse 1, D-40225 Düsseldorf, Germany; Interdisciplinary Research Center HALOmem, Charles Tanford Protein Center, Martin Luther University Halle-Wittenberg, Kurt-Mothes-Straße 3a, D-06120 Halle (Saale), Germany; UniversitätsSpital Zürich, Rämistrasse 100, CH-8091 Zürich, Switzerland; Institute of Biochemistry and Biotechnology, Martin-Luther University Halle-Wittenberg, Kurt-Mothes-Straße 3, D-06120 Halle (Saale), Germany; Biozentrum, Martin Luther University Halle-Wittenberg, Weinbergweg 22, Halle/Saale, Germany

**Author notes:** Corresponding Author: Luz Irina A. Calderón Villalobos, Molecular Signal Processing Department, Leibniz Institute of Plant Biochemistry (IPB), Weinberg 3, D-06120 Halle (Saale), Germany.

**Keywords:** intrinsic disorder, degron, auxin, AUX/IAA, SCF^TIR1^, phytohormones, allostery

## Abstract

Cullin RING-type E3 ubiquitin ligases SCF^TIR1/AFB1-5^ and their ubiquitylation targets, AUX/IAAs, sense auxin concentrations in the nucleus. TIR1 binds a surface-exposed degron in AUX/IAAs promoting their ubiquitylation and rapid auxin-regulated proteasomal degradation. Here, we resolved TIR1·auxin·IAA7 and TIR1·auxin·IAA12 complex topology, and show that flexible intrinsically disordered regions (IDRs) in the degron’s vicinity, cooperatively position AUX/IAAs on TIR1. The AUX/IAA PB1 interaction domain also assists in non-native contacts, affecting AUX/IAA dynamic interaction states. Our results establish a role for IDRs in modulating auxin receptor assemblies. By securing AUX/IAAs on two opposite surfaces of TIR1, IDR diversity supports locally tailored positioning for targeted ubiquitylation, and might provide conformational flexibility for adopting a multiplicity of functional states. We postulate IDRs in distinct members of the AUX/IAA family to be an adaptive signature for protein interaction and initiation region for proteasome recruitment.

## Main text

Proteolysis entails tight spatiotemporal regulation of cellular protein pools ^1, 2^. The ubiquitin-proteasome system (UPS) rules over protein turnover, and controls stimulation or attenuation of gene regulatory networks, depending on whether a degradation target is a transcriptional repressor or activator ^2^. A typical E1-E2-E3 enzymatic cascade warrants specific target ubiquitylation by catalyzing the ATP-dependent attachment of ubiquitin moieties to the target ^3^. Directly and indirectly, every single aspect of cellular integrity and adaptation is impacted by target ubiquitylation, e.g. cell cycle progression, apoptosis/survival, oxidative stress, differentiation and senescence ^4^. In SKP1/CULLIN1/F-BOX PROTEIN (SCF)-type E3 ubiquitin ligases, the interchangeable F-BOX PROTEIN (FBP) determines specificity to the E3 through direct physical interactions with the degradation target ^5, 6^. UPS targets carry a short degradation signal or degron, located mostly within structurally disordered regions, which is precisely recognized by cognate E3 ligases ^7^. Structural disorder and conformational flexibility within UPS targets confer diversity and specificity on regulated protein ubiquitylation and degradation ^7^. Primary degrons within a protein family, whose members share the same fate, behave as islands of sequence conservation surrounded by fast divergent intrinsically disordered regions (IDRs) ^7^. Once a favorable E3-target association stage has been accomplished, one or multiple lysines residues neighboring IDRs of the target, often form a ubiquitylation zone of functional exposed ubiquitylation sites ^8–10^. Local disorder and conformational flexibility brings next an E2-loaded with Ub (E2∼Ub) into close proximity to the bound target, such that a suitable microenvironment for catalytic Ub transfer is created ^7^. Efficient degradation of UPS targets requires the 26S proteasome to bind its protein target through a polyubiquitin chain with a specific topology, and subsequently engages the protein at a flexible initiation region for unfolding and degradation ^11^. A primary degron for E3 recruitment, a ubiquitin chain, and an IDR in UPS targets build a tripartite degron, required for efficient proteasome-mediated degradation ^7^.

Intrinsically disordered proteins (IDPs) often function in processes that underlie phenotypic plasticity such as signal transduction in plants ^12–15^. Auxin promotes plant growth and development by triggering an intracellular signaling cascade that leads to changes in gene expression ^16^. INDOLE-3-ACETIC ACID proteins (AUX/IAAs) are mostly short-lived transcriptional repressors, with half-lives varying from ∼6-80 min, and whose expression is largely and very rapidly (less than 15 minutes) stimulated by auxin ^17^. The *Arabidopsis* genome encodes for 29 AUX/IAAs, with 23 of them carrying a mostly conserved VGWPP-[VI]-[RG]-x(2)-R degron as recognition signal for an SCF^TIR^^1/AFB1-5^ E3 ubiquitin ligase for auxin-mediated AUX/IAA ubiquitylation and degradation ^18, 19^. Under low auxin concentrations, AUX/IAAs repress type A AUXIN RESPONSE FACTORS (ARF) transcription factors via physical heterotypic interactions through their type I/II Phox/Bem1p (PB1) domain ^19^. Once specific cells reach an intracellular auxin concentration threshold, F-BOX PROTEINS TRANSPORT INHIBITOR RESPONSE 1 (TIR1)/AUXIN SIGNALING F-BOX 1-5 (AFB1-5) increase their affinity for the AUX/IAA degron ^20, 21^. This results in an AUX/IAA ubiquitylation and degradation cycle that enables derepression of the transcriptional machinery ^22^. Since AUX/IAAs are themselves auxin regulated, once the intracellular AUX/IAA pool is replenished, they act again in a negative feedback loop repressing ARF activity ^23, 24^.

Degron-carrying AUX/IAAs and TIR1/AFB1-5 form an auxin receptor system, as auxin occupies a binding pocket in TIR1 just underneath the AUX/IAA degron ^20^. Auxin binding properties of the receptor complex are greatly determined by the specific AUX/IAA engaged in the receptor complex ^21^. Hence, different combinations of TIR1/AFBs and AUX/IAAs assemble at different auxin concentrations, allowing the sensing of fluctuating intracellular auxin concentrations ^21^.

Although we currently lack structural information on AUX/IAAs, they are postulated to adopt a modular structure according to sequence homology in different plant species *e.g.* 29, 3, 1 members in *Arabidopsis thaliana*, *Physcomitrella patens*, and *Marchantia polymorpha*, respectively ^25, 26^. Besides the primary degron motif, AUX/IAAs encompass a TOPLESS interacting motif for transcriptional repression, and the PB1 C-terminal domain (CTD) ^27^.

While the degron is absolutely necessary for AUX/IAA recruitment and degradation, it is not sufficient for full auxin binding properties of a TIR1·AUX/IAA auxin receptor pair ^21^. Intriguingly, unresolved flexible regions outside the primary degron contribute to differential co-receptor assembly ^21^, AUX/IAA destabilization ^28, 29^, basal protein accumulation ^30^, and are also decorated with specific lysine residues that undergo ubiquitylation *in vitro* ^31^.

The dynamic range of auxin sensitivity in plant cells, and by default growth and developmental responses, relies on efficient AUX/IAA processing by the UPS that we still need to mechanistically understand. The complexity of auxin signaling also underscores the importance of unveiling precisely how different AUX/IAAs can contribute to auxin sensing, engage in multifarious interactions, e.g. TIR1·AUX/IAA, AUX/IAA·AUX/IAA and AUX/IAA·ARF, and undergo ubiquitylation and degradation. While we recognize the degron as TIR1·AUX/IAA interaction motif, we lack information on how AUX/IAAs are positioned on TIR1, or whether additional structural AUX/IAA features might impact recruitment and auxin binding. Furthermore, full structure of AUX/IAA proteins has remained elusive so far, as they appear to adopt a highly flexible fold and/or form high order oligomers due to their PB1 domain.

Here, we studied the structural properties of AUX/IAAs and report on intrinsically disordered regions (IDRs) in IAA7 and IAA12 that influence TIR1·AUX/IAA interactions. We pursued a biochemical and structural proteomics approach and unveiled how flexibility in AUX/IAAs affect their conformational ensemble allowing surface accessibility of degrons. Our data demonstrate how an extended fold in AUX/IAAs is favorable for recruitment by the SCF^TIR1^, and offers a structural constraint for correct positioning on TIR1. Our data lays evidence of how AUXIAAs are fully recognized by the ubiquitylation machinery. We also offer a model of how a potential allosteric effect that fine-tunes TIR1·AUX/IAA interactions echoes into AUX/IAA-mediated control of gene expression.

## Results

### AUX/IAAs exhibit intrinsic structural disorder

Regions flanking the core GWPPVR degron motif influence AUX/IAA protein recruitment by SCF^TIR1^, impact auxin binding, and AUX/IAA degradation. A broader sequence context of the AUX/IAA degron might be therefore crucial for the adequate regulation of AUX/IAA processing and turnover, including post-translational modifications (e.g. ubiquitylation), protein-protein interactions and ligand-protein interactions ^21, 28, 29^. To probe whether structural flexibility and intrinsic disorder are common features of AUX/IAAs in general, we carried out an *in silico* analysis (IUPred2A) of the 29 members of the *Arabidopsis thaliana* AUX/IAA family (Fig. 1a, **Supplementary Figs. 1**). This allows to predict global structural disorder along AUX/IAA protein sequences, and to score the probability of disorder for every amino acid residue in a context-dependent manner ^32^. We also inspected the distribution of IDRs in AUX/IAAs outside the well-structured PB1 domain (Fig. 1a, **Supplementary Fig. 1**). We defined scores for disorder probability as high (= disordered, >0.6), intermediate (0.4-0.6), or low (= ordered, <0.4). First, IDRs occur in most AUX/IAAs and in almost all AUX/IAA subclades (Fig. 1a). IDR-located residues are enriched in the N-terminal halves of AUX/IAAs and much less so in the C-terminal PB1-domains (Fig. 1a, **Supplementary Fig. 1**). The length of the AUX/IAAs does not correlate with an enrichment of disorder segments because IAA1-4 or IAA28 (average length below 200 aa) exhibit features of disorder, while similarly small AUX/IAAs (*e.g.* IAA6, IAA15, IAA19, IAA32, or IAA34) are predicted to be well-structured. Interestingly, all non-canonical AUX/IAAs but IAA33, which lacks the core degron motif for interaction with TIR1 and auxin binding, are rather ordered. IAA33 diverged early during the evolution from the rest of the AUX/IAAs ^33^, and it belongs, together with canonical IAA26 and IAA13, to the most disordered family members. Intriguingly, AUX/IAAs such as IAA7 and IAA12, which are members of a different subclade ^19^, appear to have similar bias for IDRs (Fig. 1a, **Supplementary Fig. 1**).

**Figure 1.**
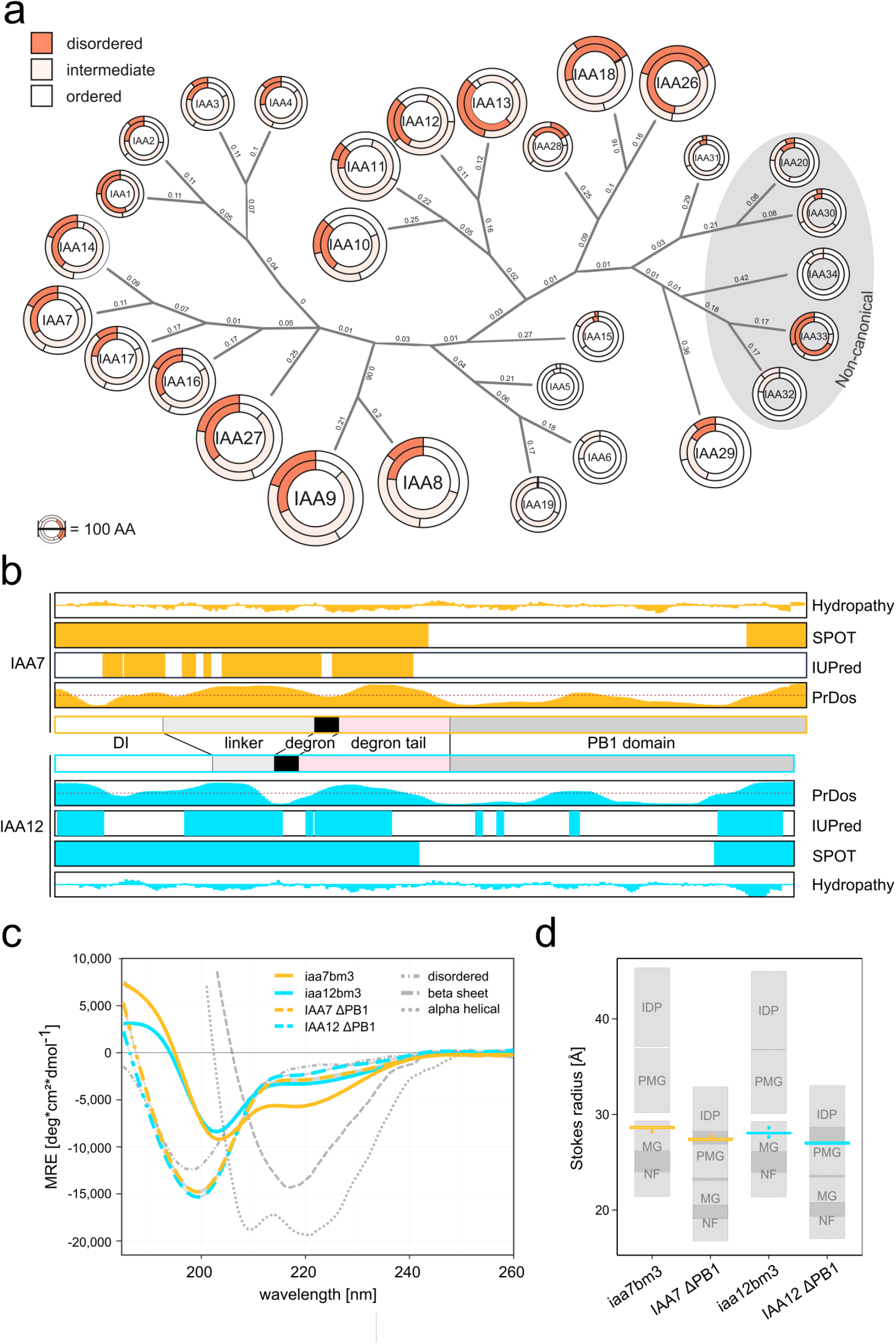
AUX/IAA proteins are intrinsically disordered outside the PB1 domain. (a) Simplified phylogenetic tree of 29 *Arabidopsis thaliana* AUX/IAAs showing their sequence composition based on IUPred2A prediction for disorder (score classification: disorder: >0.6; intermediate: 0.4-0.6; ordered: <0.4). Outer circles correspond to full length proteins, inner circles represent disorder prediction excluding the PB1 domain. (b) *In silico* prediction maps of disorder along the IAA7 and IAA12 sequence using SPOT, IUPRED1 and PrDos algorithms. AUX/IAA domain structure (Domain I (DI), a linker, a core degron, a degron tail and the Phox/Bem1p (PB1) domain) is displayed. Outer plots represent Kyte-Doolittle hydropathy (scale from −4 to +4). Dotted line in PrDos prediction represents a 0.5 threshold. (c) Circular dichroism spectra of IAA7 (orange) and IAA12 (aquamarine) oligomerization and PB1-less deficient variants (dashed colored lines) show the lack of defined secondary structure elements outside of the PB1 domain. Reference spectra (gray dotted lines) are depicted. Ellipticity is calculated as mean residual ellipticity (MRE). (d) IAA7 (orange) and IAA12 (aquamarine) exhibit an extended fold according to Stokes radii determination via size exclusion chromatography. Theoretical Stokes radii of known folds (gray, labeled rectangles): intrinsically disordered protein (IDP), pre-molten globule (PMG), molten globule (MG), natively folded (NF) plus 10% outer limits, and experimental values (colored box plots) (generated with RStudio, default settings; n = 4-9). IAA7 and IAA12 classify as MG/PMG-like proteins (bm3 variants) with less folded PMG/IDP-like features outside the PB1 domain (ΔPB1 variants).

IAA7 and IAA12 equip TIR1·AUX/IAA receptor complexes with distinct auxin binding affinities, *i.e. K*_d_ TIR1·IAA7 ∼10 nM and TIR1·IAA12 ∼ 300 nM, respectively ^21^. These differences are not exclusive to a divergent primary degron, since a TIR1-IAA12^GWPPVR^ co-receptor could not account for high auxin binding affinity ^21^. This data substantiated the hypothesis that distinct features outside the core degron, such as IDRs, might bestow AUX/IAAs, explicitly IAA7 and IAA12, with unique properties for interaction with TIR1, and therefore auxin sensitivities. In order to investigate the distribution of disorder in IAA7 and IAA12 proteins, we performed *in silico* analyses using multiple disorder prediction algorithms (Fig. 1b). Consistently, all tested algorithms showed that most of the disorder segments in IAA7 and IAA12 are located on their N-terminal half (upstream of the PB1), and near their C-terminus resembling probably a disordered “piggy tail”. We also observed an enrichment of hydrophilic residues in these IDRs (hydropathy index), which may therefore be solvent exposed (Fig. 1b). Disorder in IAA7 and IAA12 represents almost 50% of their amino acid content. In IAA7 but most notably in IAA12, we observed a predominant “order-dip” corresponding to the core degron (Fig. 1b).

Using recombinantly expressed proteins, we further analyzed IAA7 and IAA12 secondary structure and overall shape via CD spectroscopy and size exclusion chromatography, respectively (Fig. 1c-d, **Supplementary Figs. 2-3)**. Hereby, we addressed a function-related transient AUX/IAA fold while considering different protein conformational classes. In addition to wild-type and oligomerization-deficient IAA7 and IAA12 full-length proteins (iaa7bm3, iaa12bm3), we incorporated truncated variants of IAA7 and IAA12 lacking the compact PB1 domain. Both iaa7bm3 and iaa12bm3 exhibit a rather complex mix of secondary structure elements characteristic of molten globule–like proteins, displaying a minimum at ∼205 nm, and a shoulder near 220 nm in CD spectra ^34^. CD spectra of PB1-lacking IAA7 and IAA12 appear to be shifted toward a shorter wavelength with a minimum at just below 200 nm, which is characteristic for random-coil proteins (Fig. 1c, **Supplementary Fig. 2**). Figure 1d shows the measured Stokes radii (R_S_) for iaa7bm3, iaa12bm3 together with the theoretical values of IAA7 and IAA12 displaying specific folds, *v.z.* native fold (NF), molten globular (MG), premolten globule (PMG), and unfolded (IDP). Since, all measured Stokes radii are larger than the ones expected for their respective natively folded proteins, we concluded that iaa7bm3 and iaa12bm3 adapt extended structures mainly due to large proportions of intrinsically disordered segments outside of the compactly-folded PB1 domain.

### What is the impact of intrinsic disordered segments on auxin-dependent SCF^TIR1^·AUX/IAA associations?

IAA7 and IAA12 as well as their sister proteins IAA14 and IAA13, respectively, exhibit striking differences in their degron tail (**Supplementary Fig. 4**). While IAA7 and IAA14 have a short basic degron tail (<30 aa) linking the degron to the PB1 oligomerization domain, IAA12 and IAA13 have a longer (44 aa), highly charged (Lys, Glu, Asp) and unstructured degron tail (**Supplementary Fig. 4**). Because IAA7 and IAA12 have distinct and contrasting TIR1-interaction properties, we reasoned to pinpoint the determinants of these differences and the impact of IDRs on auxin-dependent TIR1·AUX/IAA associations by creating 16 seamless IAA7 and IAA12 chimeric proteins. We defined five different segments flanked by motifs conserved throughout the AUX/IAA family: DI (N-terminus including KR motif), core degron (VGWPP-[VI]-[RG]-x(2)-R), the PB1 domain (formerly known as DIII-DIV) and two variable IDRs connecting either the DI and degron (linker), or the degron and PB1 domain (degron tail) (Fig. 2a, **Supplementary Fig. 4**). We exchanged the modules between IAA7 and IAA12 and used the resulting chimeras for interaction assays with TIR1 in the yeast-two hybrid system (Y2H) for qualitative assessment of auxin co-receptor assembly (Fig. 2a, **Supplementary Fig. 4**). TIR1 interacts in an auxin-dependent manner with IAA7 containing all its native segments, denoted IAA(7-7-7-7-7). Expression of the ß-galactosidase reporter indicates stronger interaction of TIR1·IAA7 than TIR1·IAA12. As expected, mimicking degron mutants iaa7/axr2-1 (P87S) or iaa12/bdl (P74S) in the IAA7 or IAA12 chimeras (7-7-7m-7-7, 12-12-12m-12-12) abolished their association with TIR1 (Fig. 2a). Exchanging the disordered degron tail of IAA7 (36 aa) for the one in IAA12 (49 aa) IAA(7-7-7-12-7) does not affect interaction with TIR1. A IAA(12-12-12-12-7-12) chimera however, associated with TIR1 much more efficiently, and in an auxin-dependent manner than wild type IAA(12-12-12-12-12). Similarly, PB1 domain exchanges between IAA7 or IAA12 affected positively the ability of the IAA(12-12-12-12-7) chimera to interact with TIR1. To investigate interdependency of the degron tail and the PB1 domain, we exchanged the flexible degron tail of IAA12 together with its corresponding PB1 domain, and fused them to IAA7 IAA(7-7-7-12-12). In this case, TIR1·IAA(7-7-7-12-12) interaction is greatly affected, while TIR1·IAA(12-12-12-7-7) interaction, although weak, remains stronger than TIR1·IAA(12-12-12-12-12) association. Next, we omitted the degron tails in either the wild type proteins or in the PB1-swapped versions. In both instances, when IAA7 lacks its native degron tail irrespective of the PB1 it carries, (7-7-7-Δ-7 or 7-7-7-Δ-12) basal interactions with TIR1 do not occur and auxin-dependent interactions are greatly diminished. On the other hand, IAA(12-12-12-Δ-12) and IAA(12-12-12-Δ-7) chimeras invariably exhibit an increased ability to interact with TIR1, independently of their expression level (**Supplementary Fig. 4**). Surprisingly, in each case when we omitted the PB1 domain of IAA7 chimeras, we observed strong interactions with TIR1. This coincidently supports previous observations where removal of the folded PB1 domain in several AUX/IAAs resulted in accelerated auxin-induced turnover ^29^. It has been postulated this effect is due to high order complexes the PB1 domains engage in, which might negatively influence AUX/IAA processing in yeast ^29^. Hence, when the PB1 domain is omitted, chimeric proteins seem to be unhindered by oligomeric interactions and readily interact with TIR1 in yeast. Of note, independently of the layout of the core degron, either GWPPVR in IAA7 or GWPPIG in IAA12, the IAA7 degron tail and PB1 combo of IAA7 favor auxin-dependent TIR1·AUX/IAA chimera interactions (**Supplementary Fig. 5**). Taking together, auxin-dependent and –independent interactions are influenced by both the degron tail and the PB1 domain, as they probably act in concert.

**Figure 2.**
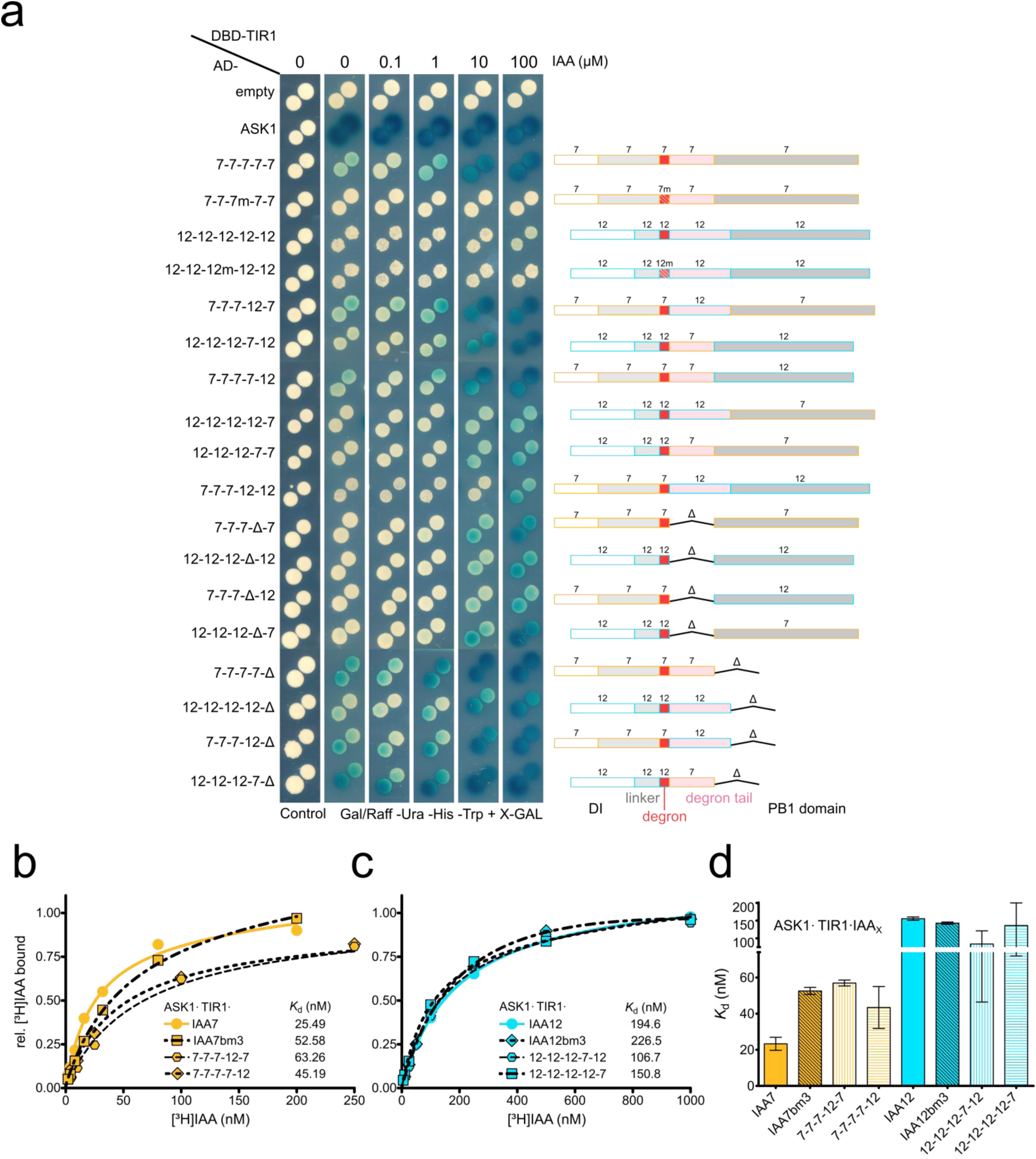
Auxin-dependent TIR1·IAA7 and TIR1·IAA12 interactions rely on the core degron, and a flexible IDR located at a suitable distance between the degron and the PB1 domain. (a) Y2H interaction matrix (left) for TIR1 with ASK1, and 16 chimeric proteins built fusing IAA7 and IAA12 segments flanked by conserved motifs throughout the AUX/IAA family. Yeast diploids containing LexA DBD-TIR1 and AD-AUX/IAA chimeras were spotted to selective medium with increasing IAA concentrations, and β-galactosidase reporter expression indicated auxin-induced TIR1-AUX/IAA interactions. AD-empty vector, negative control. Domain organization and composition of seamless chimeric IAA7 and IAA12 constructs depicted in boxes (right) with DI (white) (till KR motif), linker (light gray), core degron (red), degron tail (light pink), and PB1 domain (dark gray). Deleted domains are indicated (delta (Δ)). (b-c) Saturation binding assays using [^3^H]IAA to recombinant ASK1·TIR1·IAA7 (orange) or ASK1·TIR1·IAA12 (aquamarine) ternary complexes. TIR1·IAA7 complex exhibits a high affinity (*K*_d_ ∼20 nM) for auxin, whereas IAA12-containing co-receptor complexes provide ten-fold lower affinity for auxin (*K*_d_ ∼200 nM). Oligomerization-deficient IAA7bm3 and IAA12bm3 variants, and chimeric AUX/IAA proteins (IAA_X_) in complex with ASK1·TIR1 distinctly affect auxin bind capabilities of a co-receptor system. Shown are saturation binding curves for each co-receptor pair as relative [^3^H]IAA binding normalized to the highest value of each curve (b-c). Each point reflects technical triplicates as mean ±SEM (n=2-3). (d) Comparison of dissociation constants (*K*_d_) obtained in saturation binding experiments for each ASK1·TIR1·AUX/IAA ternary complex. Shown are mean values and standard deviation, or maximum and minimum, if applicable.

In order to address whether accessibility of IDRs and the PB1 domain in AUX/IAAs affect the outcome of TIR1·AUX/IAA interactions, we carried out *in vitro* radioligand binding assays. We used recombinant TIR1 as well as IAA7 and IAA12 chimeric or iaa7bm3 and iaa12bm3 mutant proteins. While auxin binding affinities of TIR1·iaa7bm3 and TIR1·IAA(7-7-7-7-12) complexes are reduced when compared to the TIR1·IAA7 co-receptor system (TIR1·iaa7bm3 = *K*_d_ ∼53 and TIR1·IAA(7-7-7-7-12) = *K*_d_ 45 nM vs. TIR1·IAA7 *K*_d_ ∼23 nM), we observed similar auxin affinities of TIR1·iaa12bm3 and TIR1·IAA12 co-receptors (*K*_d_ ∼195 nM vs. 226 nM, respectively) (Fig. 2b-d, **Supplementary Fig. 6**). This indicates that homotypic interactions in the case of IAA12 might not interfere with the auxin binding properties of an IAA12-containing auxin co-receptor. The decrease in the auxin binding affinity of TIR1·iaa7bm3 and TIR1·IAA(7-7-7-7-12) co-receptors hints to a positive effect of the IAA7 PB1 domain on auxin sensing (Fig. 2b and d). Exchange of disordered degron tails in chimeric IAA7 and IAA12 altered the affinity for auxin. The degron tail of IAA12 in the IAA7 context, IAA(7-7-7-12-7), reduces by two-fold auxin binding affinity of the receptor (Fig. 2b-d). Conversely, the disordered degron tail of IAA7 in an IAA12 context, IAA(12-12-12-7-12), appears to enhance AUX/IAA contribution to an auxin receptor. This is consistent with our Y2H data, where we observed a unique positive effect of the IAA7 PB1 domain. Taken together, our data confirm the postulated interdependency of the degron tail and PB1 domain, and further point to additive and separate effects of each disordered degron tail and the PB1 domain on auxin-independent and auxin-triggered TIR1 interaction.

### IDRs in AUX/IAAs harbor ubiquitylation sites or facilitate their accessibility

To investigate differences in ubiquitylation dynamics, and reveal whether the disordered nature of IAA7 and IAA12 influences their ubiquitylation rate, we recapitulated auxin-triggered and SCF^TIR1^-dependent IAA7 and IAA12 ubiquitylation ^31^. We traced IAA7 and IAA12 ubiquitylation over time and covered IAA concentrations between the auxin binding affinity (*K*_d_) of TIR1·IAA7 and TIR1·IAA12 co-receptor complexes (*i.e.* 25 nM to 155 nM) (Fig. 2d) and at higher, saturating conditions (Fig. 3, **Supplementary Fig. 7**). Ubiquitin-conjugates on IAA7 and IAA12 were traceable 10 min after incubation, and their ubiquitylation is accelerated in an auxin-dependent manner. We observed robust SCF^TIR1^-mediated IAA7 ubiquitylation even in the absence of auxin, which can be explained by previously reported basal TIR1·IAA7 interactions ^21^. IAA12∼ubiquitin conjugates were much less abundant than IAA7 after 30 min incubation (Fig. 3a). Once an auxin concentration above the *K*_d_ for the TIR1·IAA12 (±150 nM) co-receptor complex is reached however, the differences in ubiquitin conjugation on IAA7 and IAA12 are negligible over time. Reaching an auxin concentration of 1 µM corresponding to at least 50-times the *K*_d_ of TIR1·IAA7, and 4-5-times the *K*_d_ of TIR1·IAA12 for IAA (Fig. 2d), AUX/IAA ubiquitylation becomes solely time-dependent. We conclude IAA7 and IAA12 ubiquitylation occurs rapidly, and the differences in ubiquitylation dynamics depend on the auxin binding affinity by their corresponding receptors when in complex with TIR1. Given auxin to be a “molecular glue” between TIR1·AUX/IAAs, we postulate auxin might be needed for increasing the dwell-time of flexible AUX/IAAs on TIR1. This facilitates sampling of favorable AUX/IAA conformations allowing efficient ubiquitin transfer to lysine residues.

**Figure 3.**
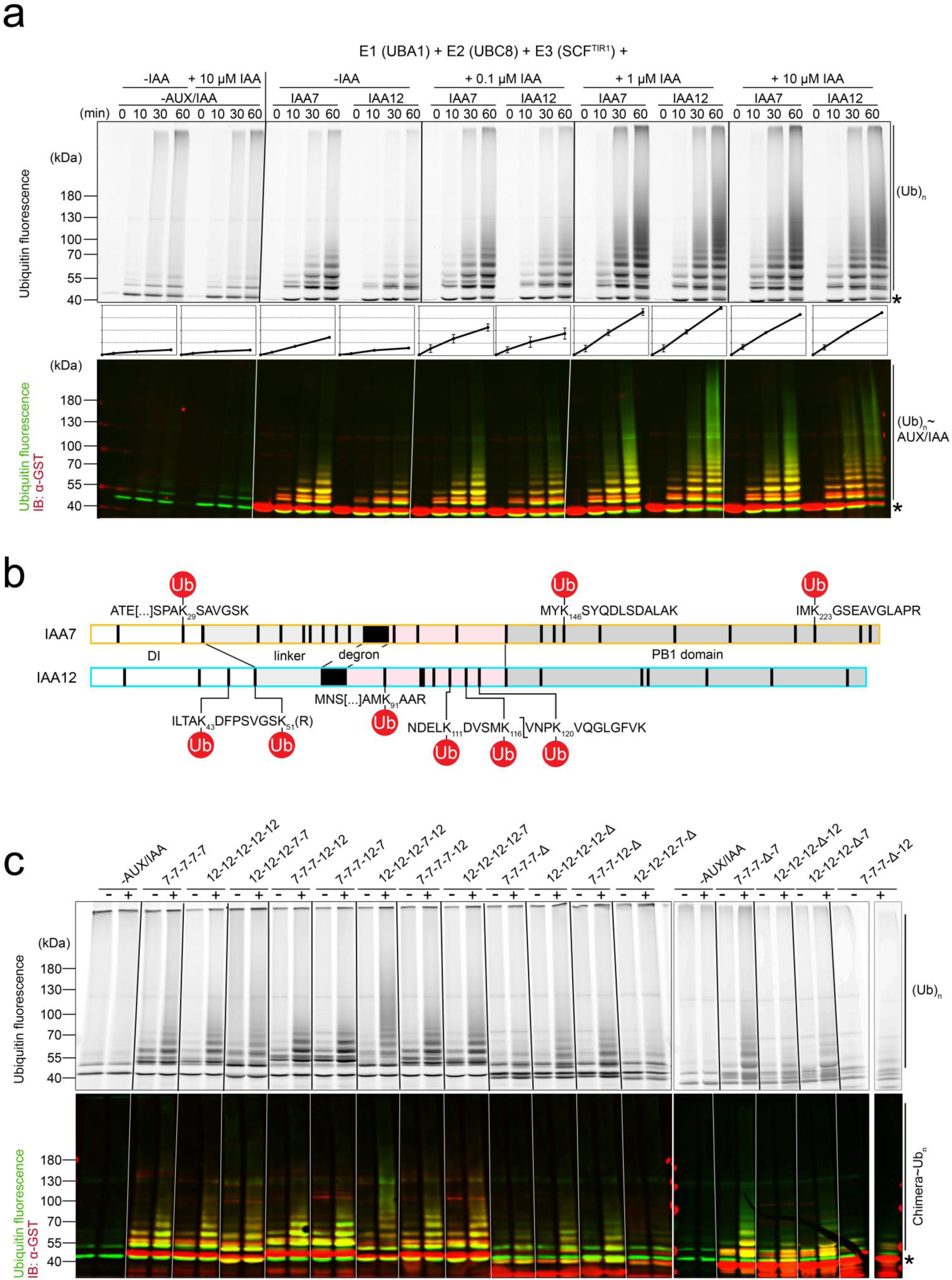
Auxin-driven and SCF^TIR1^-dependent ubiquitylation of IAA7 and IAA12 display distinct dynamics. (**a**) IVU assays with recombinant GST-IAA7 or GST-IAA12, E1 (AtUBA1), E2 (AtUBC8), reconstituted SCF^TIR1^ (*At*SKP1·TIR1, *Hs*Cul1 and *Mm*RBX1), fluorescein-labeled ubiquitin (Ub) and IAA (auxin). IAA7 and IAA12 ubiquitylation is auxin-driven and time-dependent. Basal ubiquitylation (auxin-independent) of IAA7 starts in less than 30 mins. Ubiquitylation was monitored using the ubiquitin fluorescent signal (green), and anti-GST/Alexa Fluor 647-conjugated antibodies for detection of GST-AUX/IAAs (red). ImageQuantTL software was used for quantification (middle), and generation of merged image (bottom). (**b**) IAA7 and IAA12 IVU samples were analyzed via LC-MS, and putative ubiquitylation sites detected by the diGly (or LRGG) Ub remnant after tryptic digest, were mapped relative to the domain structure. IAA12 Ub sites agglomerate in the region upstream of the degron (white) and the degron tail (light pink). (**c**) Ubiquitin-conjugation on chimeric IAA7 and IAA12 proteins in the presence or absence of 1 µM IAA. IVU reaction time 1h. Ubiquitin conjugates on chimeric proteins lacking a degron tail or the PB1 domain are evidently reduced, which is in agreement with the identified IAA7 and IAA12 Ub sites. (*) Asterisks depict the unmodified AUX/IAAs.

Putative ubiquitin acceptor lysine residues along the IAA7 and IAA12 sequences are enriched in the degron tail of IAA12, and the linker of IAA7, both of which appear to lack a three dimensional (3D) structure (Fig. 3b). We aimed therefore at gaining experimental evidence of IAA7 and IAA12 ubiquitylation sites, after *in vitro* ubiquitylation (IVU) reactions, tryptic digest and LC/MS analysis. We were able to map only few specific lysine residues on IAA7 and IAA12, which are differently distributed along their sequence (Fig. 3b, **Supplementary Table 1**). Although IAA7 and IAA12 contain 24 and 18 lysine residues, respectively, only 3 and 6 of them were ubiquitylated. While we observed only few ubiquitylated lysine residues at the AUX/IAA N-terminus, most of the mapped ubiquitylation sites were located in the region downstream of the degron, either in the PB1 domain in IAA7, or the degron tail in IAA12. Even though 4 lysines are conserved in the PB1 domain of IAA7 and IAA12, only the non-conserved residues appeared to be ubiquitylated in IAA7. The flexible degron tail of IAA7 did not get ubiquitylated, whereas 4 out of 7 lysine residues in the slightly longer disordered IAA12 degron tail could be mapped as ubiquitylation sites (Fig. 3b, **Supplementary Table 1**).

To further investigate whether the apparent structural divergence of IAA7 and IAA12 imposes restrictions to lysine access for ubiquitylation, we used chimeric IAA7 and IAA12 proteins (Fig. 3c) in our IVU assay. As we aimed at visualizing absolute differences in ubiquitin conjugation, we traced auxin-dependent ubiquitin conjugation of chimeric AUX/IAAs at a fixed IAA concentration of 0.5 µM after 1 hour IVU reaction. Exchanging the degron tails or the PB1 domains between IAA7 and IAA12 led to differences in ubiquitylation profiles of chimeric proteins compared to their wild type counterparts. This happens as we either added or subtracted regions that contain the ubiquitin acceptor sites in the IAA7 and IAA12 chimeric proteins (Fig. 3c, **Supplementary Fig. 8**). For instance, we detected an increase of ubiquitin conjugates on IAA(7-7-7-12-7), which gains ubiquitylation sites due to the exchange of the IAA7 degron tail. Deleting the AUX/IAA degron tail or the PB1 domain in the chimeric proteins results in an overall reduction of ubiquitin conjugates on targets. Versions of IAA7 or IAA12 missing a degron tail and containing the PB1 domain of IAA12, IAA(7-7-7-Δ-12) and IAA(12-12-12-Δ-12), do not undergo auxin-triggered ubiquitylation (Fig. 3c, **Supplementary Fig. 8**). Similarly, AUX/IAA versions containing the IAA7 degron but lack a PB1 domain (IAA(7-7-7-7-Δ), IAA(12-12-12-7-Δ)) are not conjugated by ubiquitin, probably due to the loss of the mapped ubiquitin acceptor sites (Fig. 3b). Our IVU assays on AUX/IAA chimeras validate our findings showing that the IAA7 PB1 domain or the flexible IAA12 degron tail carry propitious ubiquitylation sites. Thus, we postulate AUX/IAA ubiquitylation favorably occurs in exposed regions in IAA7 and IAA12, when they are recruited by TIR1.

### TIR1·AUX/IAA ensembles are guided by the degron, but tailored by flexible degron flanking regions

Due to the relative lack of a stable 3D conformation, IDPs or proteins enriched in IDRs, such as AUX/IAAs, represent a challenge for structural biology studies. During interactions with target proteins, IDPs, particularly IDRs, may undergo conformational changes that cannot be traced easily, or captured while happening ^35, 36^. Although the *Arabidopsis* SKP1 (ASK1)·TIR1·auxin·degron crystal structure enlightened us on how auxin is perceived, we lack information on the contribution of regions flanking the AUX/IAA degron on auxin binding. Thus, without being able to structurally resolve intrinsically disordered degron flanking regions, we are hindered in our understanding of how AUX/IAA ubiquitylation targets are actually positioned on TIR1. This has evidently far-reaching implications on SCF^TIR1^ E3 ubiquitin ligase activity and ubiquitin transfer by an E2 ubiquitin conjugating enzyme.

We aimed to elucidate the driving factors for ASK1·TIR1·AUX/IAA complex assembly and to unveil how IDRs in AUX/IAAs influence positioning on TIR1. We pursued a structural proteomics approach using chemical cross-linking coupled to mass spectrometric analyses (XL-MS) (Fig. 4a). We assembled ASK1·TIR1·AUX/IAA complexes containing either IAA7bm3 or IAA12bm3 proteins in the absence or presence of auxin (IAA), and added the MS-cleavable cross-linker disuccinimidyl dibutyric urea (DSBU). Reaction products were processed for mass spectrometric analysis, which utilizes the characteristic fragmentation of DSBU to identify cross-linked residues within the AUX/IAAs and the ASK1·TIR1·AUX/IAA complex ^37–39^. Our data shows multiple intra- and inter-molecular cross-links for ASK1·TIR1 and IAA7bm3 or IAA12bm3 proteins in the presence of auxin (Fig. 4b-d, **Supplementary Fig. 9-10**). In the absence of auxin, we observed only a few inter-protein and similar intra-protein cross-links when compared to auxin-containing samples (Fig. 4, **Supplementary Fig. 10**). In the presence of auxin, we identified two distinct clusters in TIR1 harboring cross-linker-reactive amino acid side chains with IAA7 and IAA12 (Fig. 4b, **Supplementary Fig. 9**). Cluster 1 comprises amino acid residues in LRR7 (217-229 aa), while cluster 2 consists of residues toward the TIR1 C-terminus located in LRR17-18 (485-529 aa). The location of the clusters on two opposing surfaces of TIR1 suggests a rather extended fold of the AUX/IAA protein when bound to TIR1 (Fig. 4b). The cross-linked residues along the sequences of ASK1·TIR1·IAA7bm3 or ASK1·TIR1·IAA12bm3 show an enrichment of highly variable intra-molecular cross-links within the AUX/IAAs (Fig. 4c-d). A low number of intra-protein cross-links along the TIR1 sequence were detected as a consequence of its rigid solenoid fold, which is in agreement with the ASK1·TIR1 crystal structure (PDB: 2P1Q, ^20^). Inter-protein cross-links indicate that the cross-linker-reactive clusters in TIR1 mainly connect with only a specific subset of AUX/IAA residues (Fig. 4b). Multiple IAA7 residues upstream of the core degron, including the KR motif, preferably cross-linked to TIR1 cluster 2. While residues downstream of the core degron, including the PB1 domain, positioned towards TIR1 cluster 1 (Fig. 4c). IAA12 is similarly positioned on TIR1, but exhibits even higher flexibility given the more diverse distribution of inter-protein cross-links (Fig. 4d). This is also supported by the fact that we detected many more assemblies for ASK1·TIR1·IAA12bm3 across replicates, than for the ASK1·TIR1·IAA7bm3 complex (Fig. 4c-d). In conclusion, our structural proteomics approach confirmed AUX/IAAs IAA7 and IAA12 exhibit flexible conformations in solution (intra-protein cross-links), and adopt an extended fold when bound to TIR1.

**Figure 4.**
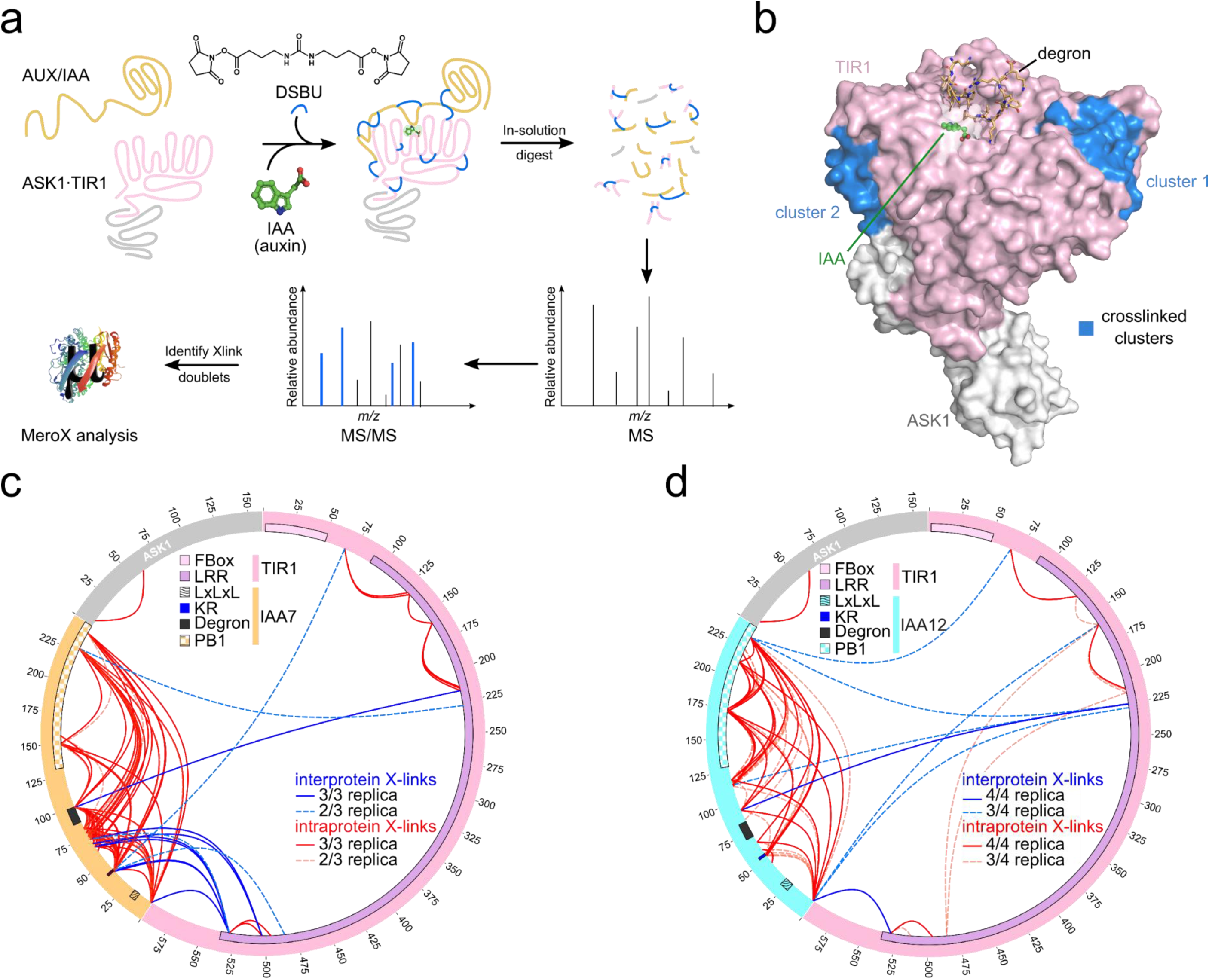
Structural proteomics using an MS-cleavable cross-linker reveals TIR1·IAA7 and TIR1·IAA12 interaction interfaces. (**a**) Workflow for the cross-linking coupled to mass spectrometry (XL-MS) approach. Recombinant oligomeric-deficient IAA7 (orange) and IAA12 (aquamarine) proteins, and ASK1·TIR1 (gray and light pink) were incubated with the DSBU cross-linker, and samples were analyzed using LC/MS/MS. Cross-linked peptides were identified using the MeroX software. (**b**) Interaction interfaces (blue) on TIR1 converge in two distinct patches around residues K217, K226 and T229 (cluster 1) or K485, S503 and K529 (cluster 2) revealing AUX/IAAs adopt an extended fold when in complex with TIR1. (**c-d**) Circular depiction of inter-protein (blue) and intra-protein (red) cross-links along IAA7 (orange), IAA12 (aquamarine), TIR1 (light pink) and ASK1 (gray) protein sequence. Cross-links were identified in at least 2/3 or 3/4 independent experiments (dashed: 2/3 and 3/4; solid lines: 3/3 and 4/4). High number of intra-protein cross-links (red) within AUX/IAAs show a high degree of flexibility characteristic of intrinsically disordered regions (IDRs). Specific cross-links within TIR1 are in agreement with the crystal structure (PDB: 2P1Q). Inter-protein crosslinks (blue) in the ASK1·TIR1·IAA7 complex mainly occurred between the N-terminus of IAA7 and the C-terminus of TIR1. Regions downstream of the IAA7 and IAA12 degron preferably cross-linked with a distinct region on TIR1 (K217, K226, T229). Known motifs and protein domains are displayed. High variability in TIR1·IAA12 cross-links hints toward a flexible and less-defined interaction interface, which reflect on the low auxin binding affinity of a TIR1·IAA12 co-receptor complex.

As we gained a better understanding on the extended fold of IAA7 and IAA12 on TIR1, we wondered whether intrinsic disordered stretches flanking the degron might help to coordinate positioning of the folded PB1 domain. An extended AUX/IAA configuration on TIR1 would be particularly relevant for allowing K146 and K223 in the PB1 domain of IAA7 to be readily available for ubiquitylation. In the case of IAA12, an assertive extension of the degron tail would expose K91, K111, K116 and K120 for ubiquitin attachment (Fig. 3b).

### Conformational heterogeneity in flexible IDR steers AUX/IAA molecular interactions

To further investigate how the intrinsic disorder in IAA7 and IAA12 influence their positioning on ASK1·TIR1, we combined our cross-linking information with a molecular docking strategy (Fig. 5, **Supplementary Fig. 11**). For that, we use available structures for the PB1 domains of AUX/IAAs and ARFs ^40–43^. We docked homology-modeled PB1 domains of *Arabidopsis* IAA7 and IAA12 to the ASK1·TIR1 complex, applying distance restraints based on the cross-linking data (Fig. 5, **Supplementary Fig. 11**). We also added an additional distance restraint reflecting the possible conformational space covered by the respective degron tails. We visualized the impact of the different restraints on the possible interaction interface of ASK1·TIR1·IAA7^PB1^ and ASK1·TIR1·IAA12^PB1^ by DisVis ^44^ (Fig. 5c-d). Evidently, by incorporating more distance restraints, we limit the number of ASK1·TIR1·AUX/IAA^PB1^ protein complexes, therefore reducing their explored interaction space (Fig. 5).

**Figure 5.**
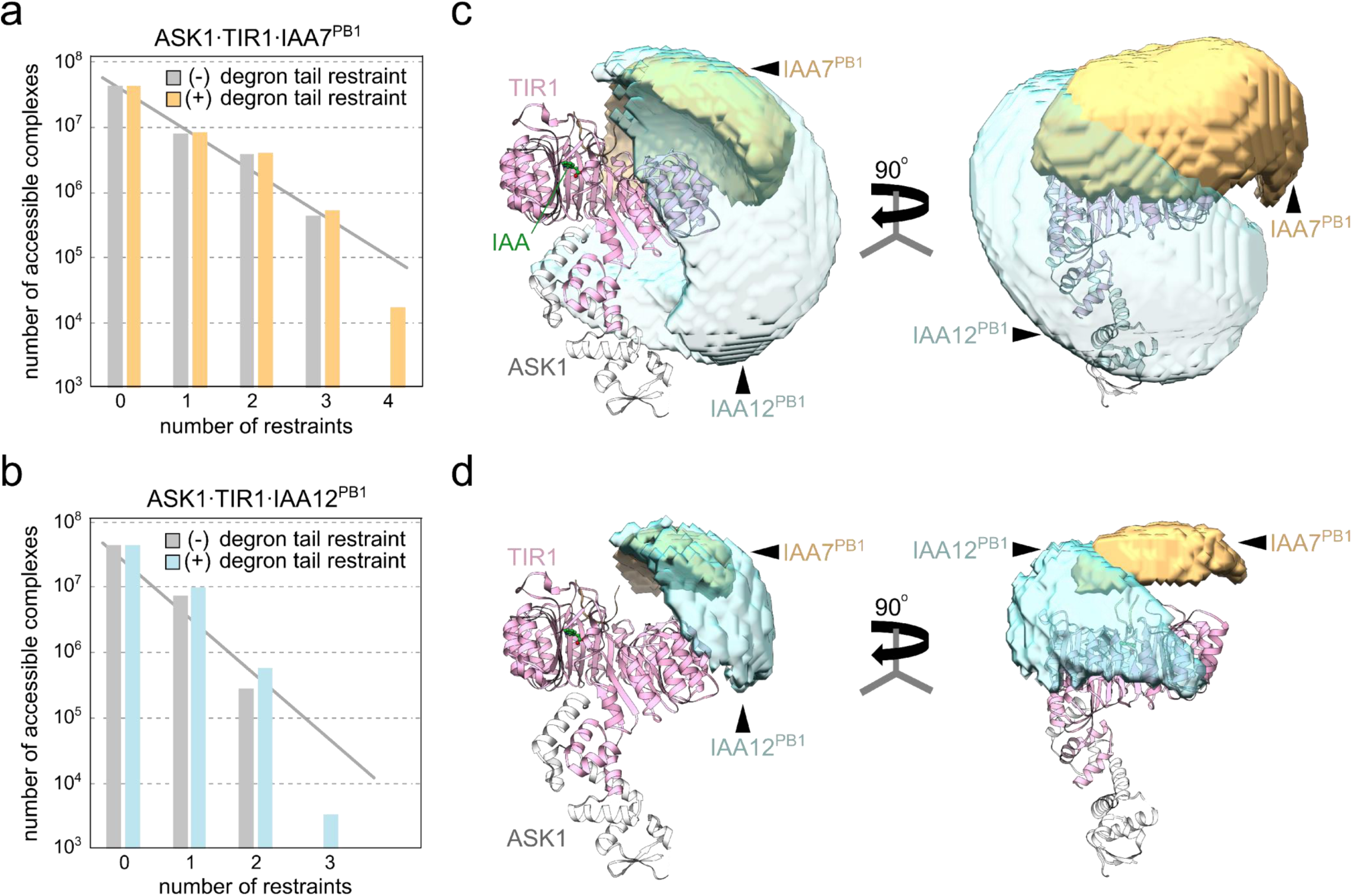
Cross-linking-based docking substantiates the function of the disordered degron tail positioning the PB1 domain of AUX/IAAs on TIR1. PB1 domains of IAA7 (light orange) and IAA12 (aquamarine) were docked on the ASK1·TIR1 (gray, light pink) structure via HADDOCK using cross-linking data as main constraint. (a-b) Including the length of the disordered degron tail of IAA7 (36 aa) or IAA12 (49 aa) as additional restraint, conspicuously reduces the conformational space, and the number of accessible TIR1·AUX/IAA PB1 complexes (c-d) Visualization of the possible conformational space occupied by the PB1 domain on the ASK1·TIR1 protein complex without (c) or with (d) the degron tail length as distance restraint. The possible interaction space of the IAA12 PB1 domain on TIR1 is much broader than the IAA7 PB1. This is an evidence the conformational space for the IAA7 PB1 on TIR1 is much more restricted, especially when the degron tail restraint is built-in. See **Supplementary Fig. 12** for best-scoring atomic detailed models.

Intriguingly, the relationship between the number of accessible complexes *vs*. the number of restraints applied does not reveal a linear behavior, but shows a sharp drop when the degron tail restraint is added to all cross-link-based restraints (Fig. 5a-b). Comparing the groups of water-refined HADDOCK models lead to similar observations and the best scoring groups were only sampled incorporating the degron tail restraint (**Supplementary Table 2**). This indicates the disordered degron tail restricts the conformational space explored by the PB1 domain (**Supplementary Table 2**, **Supplementary Fig. 11-12**). The reduction of accessible ASK1·TIR1·IAA7^PB1^ and ASK1·TIR1·IAA12^PB1^ complexes for docking is also reflected by the decreased space that can be possibly occupied by the PB1 domain (Fig. 5c-d). Overall, cross-linking-based docking of the PB1 domain of IAA12 on the ASK1·TIR1 complex is less-defined, and occupies a distinct conformational space than the ASK1·TIR1·IAA7^PB1^ complex.

In order to refine our docking data and identify the most energetically-favored TIR1·AUX/IAA^PB1^ assemblies, we carried out molecular dynamic simulations coupled to free-binding energy calculations by MM/GBSA. We used as a starting structure (t=0) the results from the HADDOCK simulations including the degron tail restraint, and performed 20 ns simulations for each TIR1·IAA7^PB1^ or TIR1·IAA12^PB1^ complex (Fig. 6 a-b). We obtained the effective binding free energy every 1 ps for each simulation, and observed distinct average effective energy (ΔG_eff_) for the different groups in each system (protein complex). Group 1 for TIR1·IAA7^PB1^ and groups 1 and 3 for TIR1·IAA12^PB1^ turned out to be energetically less favored, while groups 2 in each case showed the lowest binding energy. This indicates groups 2 likely depict the most probable ensembles (Fig. 6 a-b). To identify relevant residues in groups 2 favoring TIR1·AUX/AA interactions, we carried out per-residue effective energy decomposition analysis (prEFED) followed by validation via computational alanine scanning (CAS) (Fig. 6c, **Supplementary Table 3**). We found residues in TIR1 that might engage in polar interactions with the AUX/IAA PB1 domain. D119, D170, V171, S172, H174, H178, S199, R220 along the LRR3-6 in TIR1 likely contribute to stabilization of the TIR1·IAA7 PB1 complex. Residues H174, H178, S199 also stabilized TIR1·IAA12 PB1 interactions together with R156, S177, S201, and R205 in TIR1 LRR4-6 (Fig. 6c-d, **Supplementary Fig. 13**).

**Figure 6.**
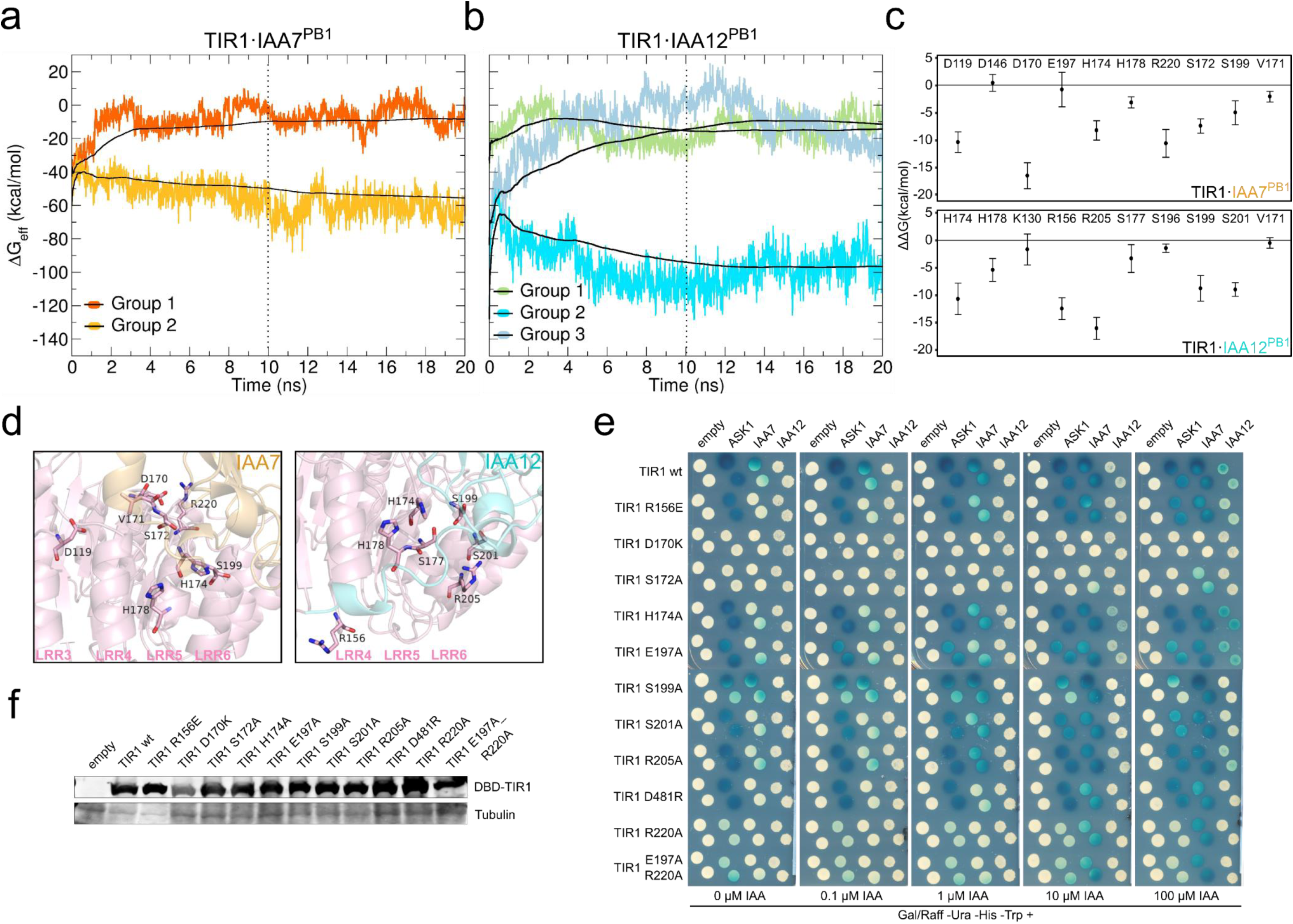
Molecular dynamics (MD) simulations using HADDOCK-based docking models reveal energetic favorable TIR1·AUX/IAA PB1 interacting moieties. (a-b) Time evolution of instantaneous Δ*G*_eff_ values over 20 ns identifying and refining stable TIR1·AUX/IAA PB1 complexes from HADDOCK best scoring groups. Black lines indicate the accumulated mean value of Δ*G*_eff_ for each trajectory. For both complexes, TIR1·IAA7 PB1 (dark and light orange) (a), and TIR1·IAA12 PB1 (blues and green) (b), one stable complex (group 2, light orange (IAA7) or aquamarine (IAA12)) characterized by a continuous low energetic state was identified. Dotted vertical line at 10 ns indicates the time point of equilibrium used as a reference for subsequent analysis. (c) Energetically relevant TIR1 residues for complex stabilization identified by computational alanine scanning (CAS) using MD trajectories (in a & b) from the equilibration time point onwards. (d) Stick representation of CAS identified residues in TIR1 (light pink) localize to the leucine-rich-repeats 3-6 (LRR3-6) forming a polar patch that allows interaction with the PB1 domain. (e) Yeast-two hybrid (Y2H) interaction matrix for TIR1 wild type and TIR1-mutant versions carrying amino acid exchanges on relevant CAS-identified residues with ASK1, IAA7 and IAA12 at different auxin concentrations.

Next, we assessed empirically whether the *in silico* identified TIR1 residues contribute to TIR1·AUX/IAA interaction. We generated mutant TIR1 proteins and evaluated their interaction with either ASK1 or IAA7 and IAA12 in Y2H assays (Fig. 6e). We aimed at identifying residues in cluster 1 at the TIR1 NTD and cluster 2 in TIR1 CTD (Fig. 4b), which might provide non-native interaction interfaces with either the PB1 domain or the KR motif of AUX/IAA proteins, respectively (Fig. 6c-e, **Supplementary Table 3**). Mutations R156E, as well as S201A, and S205A either abolished or drastically impaired basal TIR1·IAA7 and auxin-driven TIR1·IAA7 and TIR1·IAA12 associations, without affecting ASK1·TIR1 assembly. This allowed us to postulate that the positioning of the PB1 domain of AUX/IAAs on a specific NTD region in TIR1 might have a favorable effect as part of the target recruitment mechanism. Specifically, IAA7 and IAA12 PB1 “piggy tails” might be in contact with R156 in TIR1. On the other hand, mutations S172A, H174A, E197A, S199A, and R220A impaired ASK1·TIR1 and TIR1·IAA7, as well as TIR1·IAA12 interactions. This data suggests these mutations causing a long range effect on TIR1 activity and probably its overall conformational stability, that is however independent on TIR1 expression levels (Fig. 6f). While D170 offered one of the best CAS scores (**Supplementary Table 3**), D170K mutation probably corresponds to a null allele, as it leads to a reduction of protein levels, and a complete disruption of TIR1 associations (Fig. 6c, e-f).

Among the mutants tested, we also included a reversed charge exchange for D481, which is located in a negative charged patch in cluster 2 of TIR1 (Fig. 4b). According to our cross-linking data, this exposed patch (*incl.* D481, S482, E459 or E506) might exert electrostatic interactions with a conserved Lys-Arg (KR) dipeptide located between the AUX/IAA N-terminus and the degron (**Supplementary Fig. 9**). The KR was previously postulated to act as auxin-responsive rate motif influencing AUX/IAA turnover, and the magnitude of this effect was correlated with the proximity of the KR to the degron ^28, 29^. Interestingly, we found evidence for the KR motif in AUX/IAAs to favor basal interactions with the CTD of TIR1, as D481R abolished TIR1·IAA7 interaction in the absence of auxin, while weakening auxin-driven TIR1·IAA7 and TIR1·IAA12 interactions (Fig. 6e).

Our interaction studies combined with a structural proteomics approach demonstrated IDRs in IAA7 and IAA12 harbor specific features that support TIR1·AUX/IAA interactions. Charged residues surrounding the KR motif, the core degron, the degron tail and the PB1 domain act in concert to secure AUX/IAA on TIR1, thereby modulating auxin binding dynamics and likely enabling efficient ubiquitin transfer.

## Discussion

Auxin is perceived by the FBP TIR1 and its ubiquitylation targets the AUX/IAA transcriptional repressors. While TIR1 adopts a compact solenoid fold, AUX/IAAs appear flexible and modular in nature as they engage in various protein interaction networks ^23, 45^. A 13-aa degron motif in AUX/IAAs seals a ligand binding groove in TIR1, securing auxin in place. To date, we lacked information on whether additional physical interactions between TIR1·AUX/IAAs influence conformation and fate. We also did not know whether additional partner interactions facilitate the formation of the final auxin receptor complex by a two-dimensional search on the part of TIR1 on the AUX/IAA surface or *vice versa*. We found IAA7 and IAA12 exhibit conformational flexibility due to the presence of IDRs along their sequence. From the TIR1·auxin·IAA7 degron structure, we observed the degron adopts a slight helical structure ^20^. Our data shows this semihelical peptide is embedded in an intrinsically disordered stretch, which represent ∼50% of the AUX/IAA sequence, and winds down in the well-folded PB1 domain.

We showed IAA7 and IAA12 have properties of a molten globule, or a loosely packed and highly dynamic conformational fold. Although IAA7 and IAA12 might exhibit conformational heterogeneity, we observed that, while in solution, they maintain unprecedented flexibility that seems to favor recruitment by the SCF^TIR1^. Computational and experimental studies have shown IDRs such as those in AUX/IAAs, act as inter-domain linkers contributing to protein-protein interactions by exclusively or partially forming binding interfaces ^15, 46, 47^. Indeed ensembles of IAA7 and IAA12 with TIR1 captured by XL-MS allowed us to visualize AUX/IAAs “kissing and embracing” TIR1 (Fig. 7). While the degron drives auxin-mediated interactions, the IDR upstream of the degron and the PB1 domain engage in transient interactions with the CTD, and the NTD of TIR1, respectively. A directional embrace of TIR1 by an open-armed AUX/IAA, strengthened by degron-flanking IDRs, is initiated by a TIR1·auxin·degron kiss or *vice versa*. This is remarkable indeed, as we show for the first time that the PB1 domain may contact the TIR1 surface.

**Figure 7.**
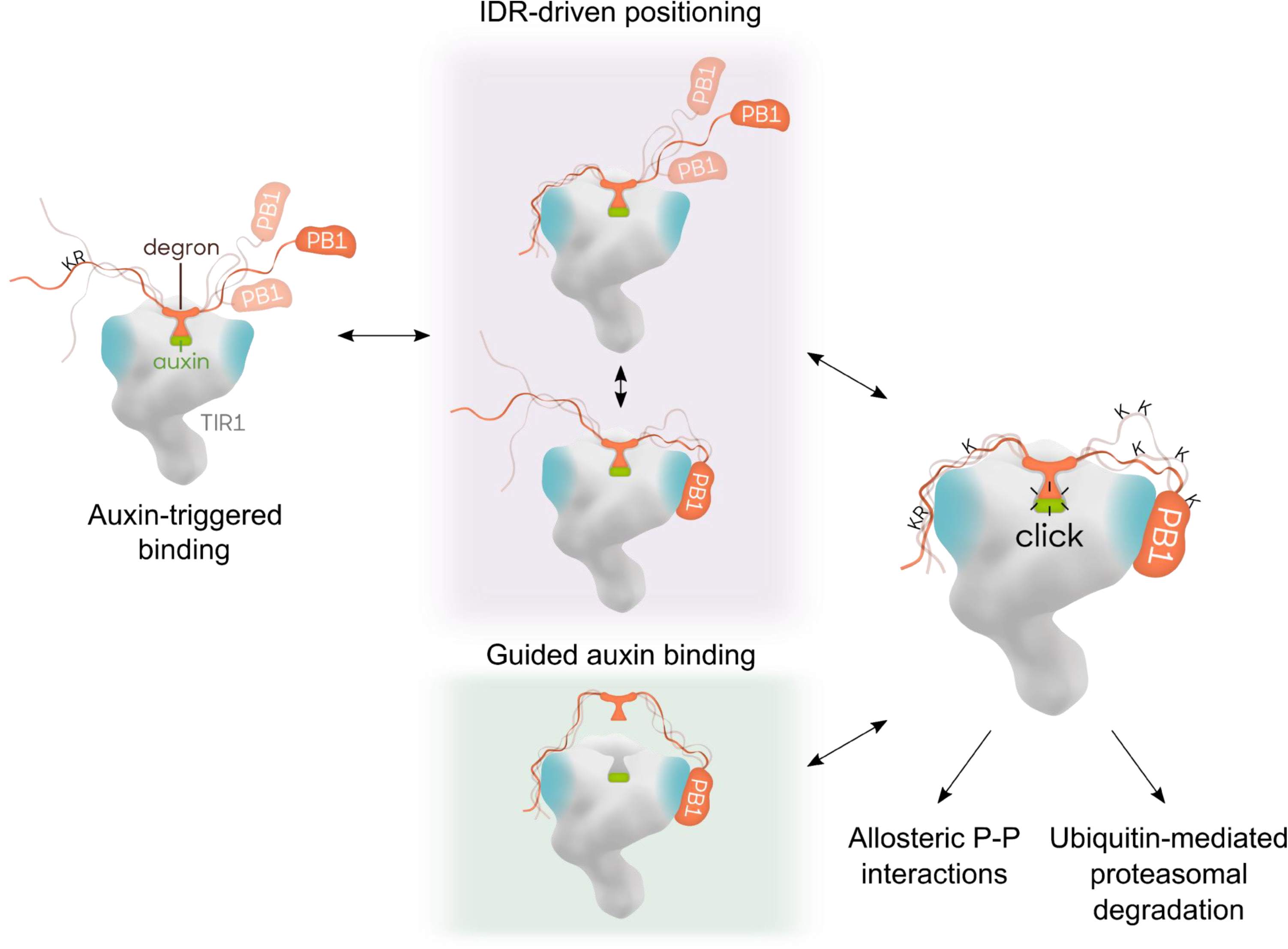
Model for ASK1·TIR1·AUX/IAA complex assembly fine-tuned by IDRs flanking the AUX/IAA core degron. The F-Box Protein TIR1 of the SCF^TIR1^ E3 ubiquitin ligase recruits AUX/IAA targets for their ubiquitylation and degradation. The phytohormone auxin and a core degron in AUX/IAAs are essential for AUX/IAA recognition. Intrinsically disordered regions (IDRs) flanking the degron provide high flexibility and an extended fold to AUX/IAAs, and influence TIR1·AUX/IAA complex formation. At least two different routes are possible for dynamic AUX/IAA recruitment and UPS-mediated degradation: *i*) auxin-triggered association between TIR1 and the core AUX/IAA degron paves the way for positioning adjacent IDRs, which exposes ubiquitin acceptor sites for efficient ubiquitylation; *ii*) transient auxin-independent interactions between IDRs, as well as the PB1 domain in AUX/IAAs and two patches of residues at opposite sides of TIR1, assist on auxin binding and offer tailored positioning. The residency time of an AUX/IAA target on TIR1, when assembled in an SCF-complex, propels processivity of AUX/IAA ubiquitylation, and impinges on availability of IDRs as initiation sites for degradation by the 26S proteasome.

Signaling proteins carrying IDRs with mostly polar and charged residues seem to have evolved more rapidly than ordered sequences, allowing increased functional complexity ^48, 49^. In our study, this applied to IDRs in regions upstream of the degron, but not for AUX/IAAs degron tails. Although IAA7 and IAA12 show differences on IDR content and length, both embraced TIR1 in a similar manner. Despite the fact that the degron tail in IAA12 is third longest in the AUX/IAA family (49 aa) and about ∼1.3 times longer than the tail of IAA7 (36 aa), both offer flexibility (**Supplementary Fig. 14**). An introduced degron length constraint in IAA7 and IAA12 greatly reduced the sampled conformational space of IAA7 and IAA12 on TIR1. Finally, degron tails seem to increase the interaction surface with TIR1, which we anticipate translates into variability of binding kinetics.

Within the *Arabidopsis* AUX/IAA protein family, nearly half of the degron tails are between 20-40 aa long and show high disorder probability (**Supplementary Fig. 1**). Seven of the 23 degron-containing AUX/IAAs (IAA19, IAA4, IAA6, IAA5, IAA1, IAA2, IAA15), however, carry a relatively ordered degron tail shorter than 20 amino acids (**Supplementary Fig. 14**). Is that specific length an evolutionary constraint for TIR1 association? Auxin-dependent gene regulation, and AUX/IAA proteins appear in the land plant lineage over 500 mya ^33, 50^. When comparing the proteins sequence of the two ancestral AUX/IAAs in moss and *Marchantia* ^25, 26^, we observed their degron tails are not much longer than the average degron tails (40 aa) of *Arabidopsis* AUX/IAAs, despite the overall length of these proteins being at least double that of angiosperm AUX/IAAs. It will be interesting to investigate whether degron tails length and disorder content are a deeply conserved features for surface availability, and whether short degron tails (less than 20 aa) can still offer tailored positioning on TIR1. Furthermore, the degron tail might generate an entropic force ^51, 52^ that is fine-tuned, but also restricted by IDR length, modulating binding of AUX/IAAs to TIR1.

Particular stretches of amino acids with increased evolutionary conservation within disordered segments have been found to determine interaction specificity, acting as functional sites ^48, 49, 53^. This seems to precisely apply to the region in AUX/IAAs upstream of the degron containing the auxin-responsive rate KR motif ^28, 54^. The KR exhibits a high level of conservation, and in addition to being part of a bipartite nuclear localization signal (NLS), the KR contributes to assembly of a TIR1·AUX/IAA auxin receptor complex and, probably as a result, is required for basal proteolysis *in planta* and AUX/IAA degradation dynamics ^21, 28, 29, 54^. How mechanistically could the KR exert an effect on TIR1 recognition and further AUX/IAA processing? Our findings lead us to propose an answer to a more than 10 year’s long standing question. As part of the AUX/IAA embrace of TIR1, the KR motif embedded in the IDR upstream of the degron offers alternative contacts with the CTD of TIR1 and probably first binding contacts (Fig. 7). Previous studies showed that moving the KR motif closer to the degron was not sufficient to accelerate AUX/IAA degradation rate ^29^. We predict a high flexibility of the IDR offers a necessary distance between the KR and the core degron for reaching distinct TIR1 contact sites. So, the positively charged KR motif in AUX/IAAs may be capable of engaging in electrostatic interactions with a cluster of highly charged (Asp, Glu) residues in TIR1 between LRR16 and LRR18, where D481 is located. While TIR1 and AFB1 offer similar contact points to the KR in AUX/IAAs, AFB2 and AFB3 exhibit opposite charged residues (Lys) that however might still provide charge-charge interactions with a specific subset of AUX/IAAs. It remains to be determined whether this is an additional feature facilitating differential auxin sensing by distinct TIR1/AFBs·AUX/IAA co-receptor combinations ^21^.

Local flexibility in AUX/IAAs is evidently shaping their conformation when in complex with TIR1. Specifically, flexible IDRs flanking the core degron in AUX/IAAs, as shown for IAA7 and IAA12, serve as variable spacers between the degron and the well-folded PB1 domain. Our data provide evidence for dynamic allosteric modulation of a TIR1·AUX/IAA auxin receptor complex by the folded-PB1 domain and IDRs in AUX/IAAs. We could track positive but also negative cooperativity, due to the degron tail and PB1 domain combo, fine-tuning conformational states of TIR1·IAA7 and TIR1·IAA12 receptor pairs, respectively. Further long-range, probable allosteric, effects are reflected into AUX/IAA turnover, when PB1 domain and degron tail act as one element (**Supplementary Fig. 5**).

Structural disorder in AUX/IAA targets appears also to be instrumental for processivity in ubiquitin transfer by the SCF^TIR1^ E3 ubiquitin ligase. This is crucial as once an active E2-E3-target assembly has formed, spatial and geometric constraints such as distance and orientation relative to the E3-bound primary degron limit ubiquitylation surface and lysine selection for degradation ^7^. AUX/IAA sequence harbors a number of putative ubiquitin acceptor lysines (∼9% total sequence) (**Supplementary Fig. 14**). Our data show that not all of these sites are favorable for ubiquitylation. Downstream of the core degron, AUX/IAAs likely offer an attractive region for ubiquitin conjugation. We predict either the PB1 or the degron tail facilitate the accessibility of receptor residues that undergo ubiquitylation. Upon TIR1·AUX/IAA interaction, IDRs either act themselves as ubiquitylation acceptor sites (e.g. IAA12) or orient the PB1 domain-located lysines as ubiquitin acceptor sites (e.g. IAA7). We cannot rule out however, regulated and efficient ubiquitin transfer might prioritize target degradation at the proteasome. This is key, as AUX/IAA turnover likely needs properly positioned ubiquitin moieties at the proper distance of an IDR, and an IDR with unbiased sequence composition as an initiation site for efficient degradation ^55–57^. To better understand this, it will be imperative to gain insights into where AUX/IAAs are ubiquitylated *in vivo*, and where exactly the proteasome initiates degradation relative to the ubiquitylation sites.

The effects of cooperative allostery driven by IDRs in AUX/IAA proteins might not be limited to the TIR1·AUX/IAA interaction, but rather influence the assembly into other complexes regulating auxin output signals ^58^. It is therefore also possible that in response to fluctuating cellular auxin concentrations, transient TIR1·AUX/IAA interactions via IDRs alter the energy landscape of AUX/IAA·TPL, AUX/IAA·ARF and AUX/IAA·AUX/IAA assemblies and/or possible decorations with PTMs. Future studies will tell whether IDRs in AUX/IAAs, and the recently described IDRs in ARFs, affect their protein assembly’s localization or activity ^59^. One can envision, IDR-driven cooperativity resulting in a multiplicity of allosterically-regulated interactions within the auxin signaling pathway, where AUX/IAAs act as signaling hubs within the different complexes.

Using an XL-MS approach, we have captured for the first time a highly flexible ubiquitylation target being engaged by an SCF-type E3 ubiquitin ligase, which at the same time, constitutes a phytohormone receptor. Our strategy offers an opportunity to visualize how IDR-driven allostery might influence a complex signaling network.

## Methods

### Phylogenetic tree generation and secondary structure analysis

Phylogenetic tree construction was done using Clustal Omega^60^ with standard settings and the full-length protein sequences of all *Arabidopsis* AUX/IAAs deposited at uniprot^61^. The constructed tree was visualized by iTOL^62^ and manually edited. *In silico* disorder analysis was performed with the web-based IUPred2A tool^32^ utilizing AUX/IAA protein sequences. The resulting disorder probability was used to categorize each residue as either ordered (<0.4), intermediate (0.4-0.6) or disordered (>0.6). Same analysis was done for all AUX/IAA proteins excluding the PB1 domain using the conserved VKV motif as the start of the PB1 domain. Residues of each category were plotted using R. IAA7 and IAA12 disorder predictions were additionally carried out using SPOT ^63^ and PrDOS ^64^ algorithms with standard settings. Hydropathy plots were generated via Expasy-linked ProtScale ^65, 66^ using the Kyte-Doolittle method ^67^.

### Protein purification

ASK1·TIR1 complex was purified from Sf9 cells as described earlier ^20^ with minor changes. In brief, ASK1 was co-purified with GST-TIR1 using GSH affinity chromatography (gravity flow) and anion chromatography (MonoQ) followed by Tag-removal and a final size-exclusion chromatography (SEC) step (Superdex 200), using an ÄKTA FPLC system.

AUX/IAA proteins, including chimeric versions, were expressed as GST-tagged proteins in *E.coli* and purified using GSH affinity chromatography, including a high salt wash (1M NaCl) and gravity flow anion exchange chromatography (Sepharose Q). For circular dichroism, the GST-tag was removed on the GSH column matrix with thrombin, and fractions containing AUX/IAAs were briefly concentrated, passed over a benzamidine column, and further purified using a Sephacryl S100 column (SEC) with an ÄKTA FPLC system. This step was carried out using the CD measurement buffer (see CD measurement section) for buffer exchange.

### Size exclusion chromatography and size calculations

The last protein purification step was used to simultaneously determine the Stokes radii of AUX/IAAs in CD buffer (10 mM KPi pH 7.8; 150 mM KF; 0.2 mM TCEP). The HiPrep 16/60 Sephacryl S-100 high resolution column was calibrated using gel filtration standards (Bio-Rad, Cat. #151-1901) with added BSA before the runs. Stokes radii for the globular known reference proteins were calculated as described^68^. The Stokes radii of AUX/IAA variants were calculated from the resulting calibration curve equation based on their retention volume (n = 4-9).

### Circular Dichroism (CD) measurements

After purification, including tag-removal and size exclusion chromatography, AUX/IAAs were concentrated and adjusted to 2.5 - 5 µM in CD buffer. CD measurements were carried out on a Jasco CD J-815 spectrometer and spectra were recorded from 260 nm to 185 nm as 32 accumulations using a 0.1 nm interval and 100 nm/min scanning speed. Cell length was 1 mm and temperature was set to 25°C. All spectra were buffer corrected using CD buffer as a control and converted to mean residual ellipticity (MRE). Reference spectra for a disordered (MG-14; PCDDBID: CD0004055000), a beta-sheet (BtuB; PCDDBID: CD0000102000) and an alpha-helical protein (amtB; PCDDBID: CD0000099000) were used.

### [3H]-labeled Auxin Binding Assay

Radioligand binding assays were performed as previously described^69^ using purified ASK1·TIR1 protein complexes, GST-tagged AUX/IAAs incl. chimeric AUX/IAAs and [^3^H]IAA with a specific activity of 25 Ci/mmol (Hartmann Analytic). Final protein concentrations in a 100 µL reaction were 0.01 µM ASK1·TIR1 complex and 0.3 µM AUX/IAAs. Complexes were allowed to form 1 h on ice, shaking. For non-specific binding controls, reactions contained additionally 2 mM cold IAA. Data was evaluated with GraphPad Prism v 5.04, and fitted using the “one site total and non-specific binding” preset.

### LexA Yeast Two Hybrid Assays

LexA-based yeast two hybrid assays were performed using yeast transformed with the described constructs (DBD-fusions: EGY48+pSH18-34, pGILDA vector; AD-fusions: YM4271, pB42AD vector), freshly mated and grown on selection medium (Gal/Raff – Ura –His –Trp). Same amount of yeast cells (OD_600_ = 0.4 or 0.8 for IAA12(-like)) were spotted on selection plates containing BU salts (final: 7 g/L Na_2_HPO_4_, 3 g/L NaH_2_PO_4_, pH 7), X-Gal (final 80 mg/L) and the given auxin (IAA) concentration. Plates were incubated at 30°C for several days and constantly monitored. Expression of chimeric AUX/IAAs and TIR1 mutants in yeast were checked using immunoblot analysis on lysates from haploid yeast. 50 mL liquid selection medium (Gal/Raff -Ura -His or -Trp) were inoculated with an 1/25 volume overnight culture and grown till OD_600_ ≈ 0.6, harvested, washed with water and lysed in 200 µL lysis buffer (0.1 M NaOH, 2 % β-mercaptoethanol, 2 % sodium dodecyl sulfate, 0.05 M EDTA, 200 µM benzamidine, 1 mM PMSF, Roche protease inhibitor cocktail) at 90°C for 10 min. After neutralization with 5 µL 4 M sodium acetate for 10 min at 90°C, 50 µL 4X Laemmli was added and samples were separated via SDS-PAGE and immunoblotted.

### In vitro reconstitution of Ub-conjugation

*In vitro* ubiquitylation (IVU) reactions were performed as previously described^31^. In brief two protein mixtures (mix A and mix B) were prepared in parallel. Mix A contained 50 µM ubiquitin (Ub; Fluorescein-labeled Ub^S20C^: Ub^K0^; 4:1 mix), 0.2 µM 6xHis-UBA1 (E1) and 2 µM 6xHis-AtUBC8 (E2) in reaction buffer (30 mM Tris-HCl, pH 8.0, 100 mM NaCl, 2 mM DTT, 5 mM MgCl_2_, 1 µM ZnCl_2_, 2 mM ATP). Mix B contained 1 µM Cul1·RBX1, 1µM ASK1·TIR1, and 5 µM AUX/IAA protein in reaction buffer. Mix B was aliquoted and supplemented with IAA to reach the indicated final concentration. Mixtures A and B were separately incubated for 5 or 10 minutes at 25 °C, respectively. Equal volumes of mix A and B were combined, aliquots were taken at specified time points, and reactions were stopped by denaturation in Laemmli buffer. IVUs with chimeric AUX/IAAs were carried out 1 h with 1 µM IAA. Immunodetection of Ub-conjugated proteins was performed using polyclonal anti-GST in rabbit (Sigma, G7781; 1:20,000) antibodies combined with secondary anti-rabbit Alexa Fluor® Plus 647 antibody (1:20,000) (Thermo Fischer Scientific, A32733). Detection was performed with a Typhoon FLA 9500 system (473 nm excitation wavelength and LPB filter for fluorescein-labeled ubiquitin signal detection and 635 nm excitation wavelength and LPR filter for GST signal).

Quantification of ubiquitylated AUX/IAAs was performed using the in-gel fluorescein signal above GST-IAA7 and GST-IAA12, as well as the ubiquitin-modified Cullin (∼50 kDa) were quantified for each lane using ImageQuant TL software automatic lane detection. The reduction of unmodified GST-IAA7 and GST-IAA12 fusion proteins was quantified after blotting and immunodetection using the Alexa Fluor 647 signal, and automatic band detection. All signals were background subtracted (rubberband method).

### LC-MS analyses of IVU reactions

Three sets of IVUs, corresponding to three biological replicates, were carried out on consecutive weeks using AUX/IAA proteins from different batch preparations. After 30 minutes, IVUs were stopped by denaturing with urea, reduced with DTT and alkylated with iodoacetamide. Trypsin digestion was carried out overnight at 37°C. Upon quenching and desalting, peptides were separated using liquid chromatography C18 reverse phase chemistry and later electrosprayed on-line into a QExactive Plus mass spectrometer (Thermo Fisher Scientific). MS/MS peptide sequencing was performed using a Top20 DDA scan strategy with HCD fragmentation. Ubiquitylated residues on identified peptides were mapped using GG and LRGG signatures (as tolerated variable modifications) from using both the Mascot software v2.5.0 (Matrix Science) linked to Proteome Discoverer v1.4 (Thermo Fisher Scientific) and the MaxQuant software v1.5.0.0. Mass spectrometry proteomics data are being currently curated at PRIDE repository, ProteomeXchange (http://www.ebi.ac.uk/pride/archive/). A dataset identifier will be shared upon acceptance of the manuscript.

### Cross-linking reactions

DSBU (ThermoFisher) cross-linking reactions were performed for 1 h at 25°C with either 4-5 µM of ASK1·TIR1 and 5 µM iaa7bm3 or iaa12bm3 or 10 µM iaa7bm3 or iaa12bm3 alone. Proteins were pre-incubated 15 minutes in the presence or absence of 10 µM auxin (IAA) before addition of 1 mM DSBU (100 molar excess). After TRIS quenching, samples were sonicated in the presence of sodium deoxycholate, reduced with DTT, and alkylated with iodoacetamide. Alkylation was quenched by DTT, and the reactions were incubated with trypsin over night at 37°C. Digestion was stopped with 10% TFA and after centrifugation (5 min 14.000 xg), peptide mixtures were analyzed via LC/MS.

### Mass spectrometry analyses of cross-linked peptides & data analysis

Proteolytic peptide mixtures were analyzed by LC/MS/MS on an UltiMate 3000 RSLC nano-HPLC system coupled to an Orbitrap Fusion Tribrid mass spectrometer (Thermo Fisher Scientific). Peptides were separated on reversed phase C18 columns (trapping column: Acclaim PepMap 100, 300 μm × 5 mm, 5μm, 100 Å (Thermo Fisher Scientific); separation column: self-packed Picofrit nanospray C18 column, 75 μM × 250 mm, 1.9 μm, 80 Å, tip ID 10 µm (New Objective)). After desalting the samples on the trapping column, peptides were eluted and separated using a linear gradient from 3% to 40% B (solvent A: 0.1% (v/v) formic acid in water, solvent B: 0.08% (v/v) formic acid in acetonitrile) with a constant flow rate of 300 nl/min over 90 min. Data were acquired in data-dependent MS/MS mode with stepped higher-energy collision-induced dissociation (HCD) and normalized collision energies of 27%, 30%, and 33%. Each high-resolution full scan (m/z 299 to 1799, R = 120,000 at m/z 200) in the orbitrap was followed by high-resolution product ion scans (R = 30,000), starting with the most intense signal in the full-scan mass spectrum (isolation window 2 Th); the target value of the automated gain control was set to 3,000,000 (MS) and 250,000 (MS/MS), maximum accumulation times were set to 50 ms (MS) and 200 ms (MS/MS) and the maximum cycle time was 5 s. Precursor ions with charge states <3+ and >7+ or were excluded from fragmentation. Dynamic exclusion was enabled (duration 60 seconds, window 2 ppm).

For cross-linking analysis, mass spectrometric *.raw files were converted to mzML using Proteome Discoverer 2.0. MeroX analysis was performed with the following settings: Proteolytic cleavage: C-terminal at Lys and Arg with 3 missed cleavages, peptides’ length: 5 to 30, static modification: alkylation of Cys by IAA, variable modification: oxidation of M, cross-linker: DSBU with specificity towards Lys, Ser, Thr, Tyr, and N-termini, analysis mode: RISE-UP mode, minimum peptide score: 10, precursor mass accuracy: 3 ppm, product ion mass accuracy: 6 ppm (performing mass recalibration, average of deviations), signal-to-noise ratio: 1.5, precursor mass correction activated, prescore cut-off at 10% intensity, FDR cut-off: 1%, and minimum score cut-off: 60. For further analysis only cross-links found in at least 2/3 (IAA7) or 3/4 (IAA12) experiments were considered.

### Cross-link-based docking using HADDOCK

Comparative models of IAA7 and IAA12 PB1 domains were created using multi-sequence-structure-alignments (PIR formatted) as input for MODELLER 0.921 ^70^. The generated models were incorporated for the HADDOCK-based docking together with the available ASK1·TIR1 structure (PDB code: 2P1Q, resolution: R= 1.91 Å)^20^. A detailed description how to prepare pdb files and incorporated distance restraint can be found elsewhere. Formatted pdb files were uploaded to the HADDOCK server ^71, 72^ using guru access level. To incorporate distance restraints, we used distances from intramolecular cross-links of known distance (see SUPP file docking parameters). We further added a distance restraint or not corresponding to the degron tail length calculated as described ^73^. For each complex docked, 10,000 rigid body docking structures were generated followed by a second iteration (400 best structures). Finally, 200 models/structures were water refined (explicit solvent) and clustered (FCC^74^ at 0.6 RMSD cutoff).

Using the same restraints, the possible conformational docking space of the PB1 domains was searched and visualized using DisVis^44, 75, 76^ with standard parameters. For image creation PyMOL^TM^ (Version 2.1) and UCSF Chimera^77^ were used.

### Molecular dynamic simulations (MDS) of protein-protein complexes

One refined structure of each group, derived from the cross-link-based docking by HADDOCK incorporating the disorder restraint (2 groups for IAA7^PB1^·TIR1; 3 groups for IAA12^PB1^·TIR1), was used as starting structure for MD simulations. The 5 structures were prepared using structure preparation and protonate 3D (pH = 7.5) modules and subsequently minimized with AMBER10 force-field in MOE 2019.0101 (Chemical Computing Group Inc., Montreal, Quebec, Canada).

Molecular dynamic simulations were performed with the GROMACS software package (version 4.6.5)^78^. The parameters corresponding to the proteins were generated with AMBER99SB-ILDN force-field^79^, TIP3P explicit solvation model^80^ was used and electro-neutrality was guaranteed with a NaCl concentration of 0.2 mol/L. The protocol employed here to perform MD simulations involves prior energy minimization and position-restrained equilibration, as outlined by Lindahl ^81^ for lysozyme in water. Once the system was equilibrated, we proceeded to the productive dynamic simulation without position restraint for 20 ns.

### Effective binding free energy calculations using MM-GBSA

The effective binding free energy (Δ*G_eff_*) of the protein-protein complexes formation was calculated using MMPBSA.py from Amber18 package employing the MM-GBSA method ^82^. We followed the single trajectory approach, in which the trajectories for the free proteins were extracted from that of the protein-protein complexes. *GB^OBC1^* and *GB^OBC2^* implicit solvation models were employed^82^. The accumulated mean value of Δ*G_eff_* were obtained every 10 ps from the productive MD simulation.

Energetically-relevant residues (hot-spots) at the interfaces of TIR1·AUX/IAA PB1 complexes were predicted by using the per-residue effective free energy decomposition (prEFED) protocol implemented in MMPBSA.py ^82^. Hot-spot residues were defined as those with a side-chain energy contribution (Δ*G_sc_*) of ≤ −1.0 kcal/mol. We used Computational Alanine Scanning (CAS)^82^ to further assess per-residue free energy contributions. Alanine single-point mutations were generated on previously identified hot-spots from the prEFED protocol. Both prEFED and CAS protocols were performed from the last 10 ns of the MD simulation.

## Supporting information

Supplementary Information_Niemeyer et al

Niemeyer et al_Supp Table 1

## Acknowledgements

We thank Wolfgang Brandt for initial in silico models of AUX/IAA PB1 domains, and Silvestre Marillonet for the design of constructs for Golden Gate Technology. Thanks to Steffen Abel and Elisabeth Chapman for providing input to the manuscript. This work was supported by the Deutsche Forschungsgemeinschaft (DFG, research project CA716/2-1), and core funding of the Leibniz Institute of Plant Biochemistry (IPB).

## Author contributions

M.N. and L.I.A.C.V. prepared the manuscript and designed experiments. M.N. performed biochemical experiments and analyzed the data. M.N., C.I. and C.H.I. carried out XL-MS experiments and data analysis. P.K. and M.N carried out all HADDOCK-based approaches including DisVis, and E.M.C. computational calculation and simulations. M.N., A.H. and V.W. generated Y2H constructs and performed the assays. M.N. performed ubiquitylation experiments, and together with W.H. analyzed mass spectral data of ubiquitylation sites. S.S. and M.Z. carried out ratiometric experiments and analyzed the data. E.M.C., C.I, C.I., P.K., and A.S. provided input to the manuscript. All authors approved the intellectual content.

## Competing Interest Statement

The authors declared no competing interests.

## Additional Information

Supplementary information is available online. Mass spectrometry proteomics data have been deposited to the ProteomeXchange Consortium via the PRIDE partner repository with the data sets identifiers: PXD015285 (XL-MS) and PXD015392 (ubiquitylation site identification data). Correspondence and request for materials should be addressed to L.I.A.C.V.

## Notes

#### Summary of Updates

Title updated; the main text and methods section were streamlined and shortened; main Figure 2 and Acknowledgements have been updated. Set identifiers corresponding to the Mass spectrometry proteomics data deposited to the ProteomeXchange Consortium (PRIDE repository) have been brought up to date.

