## Supplementary Information_Niemeyer et al for "Flexibility of intrinsically disordered degrons in AUX/IAA proteins reinforces auxin co-receptor assemblies"

### Supp. Fig. 1

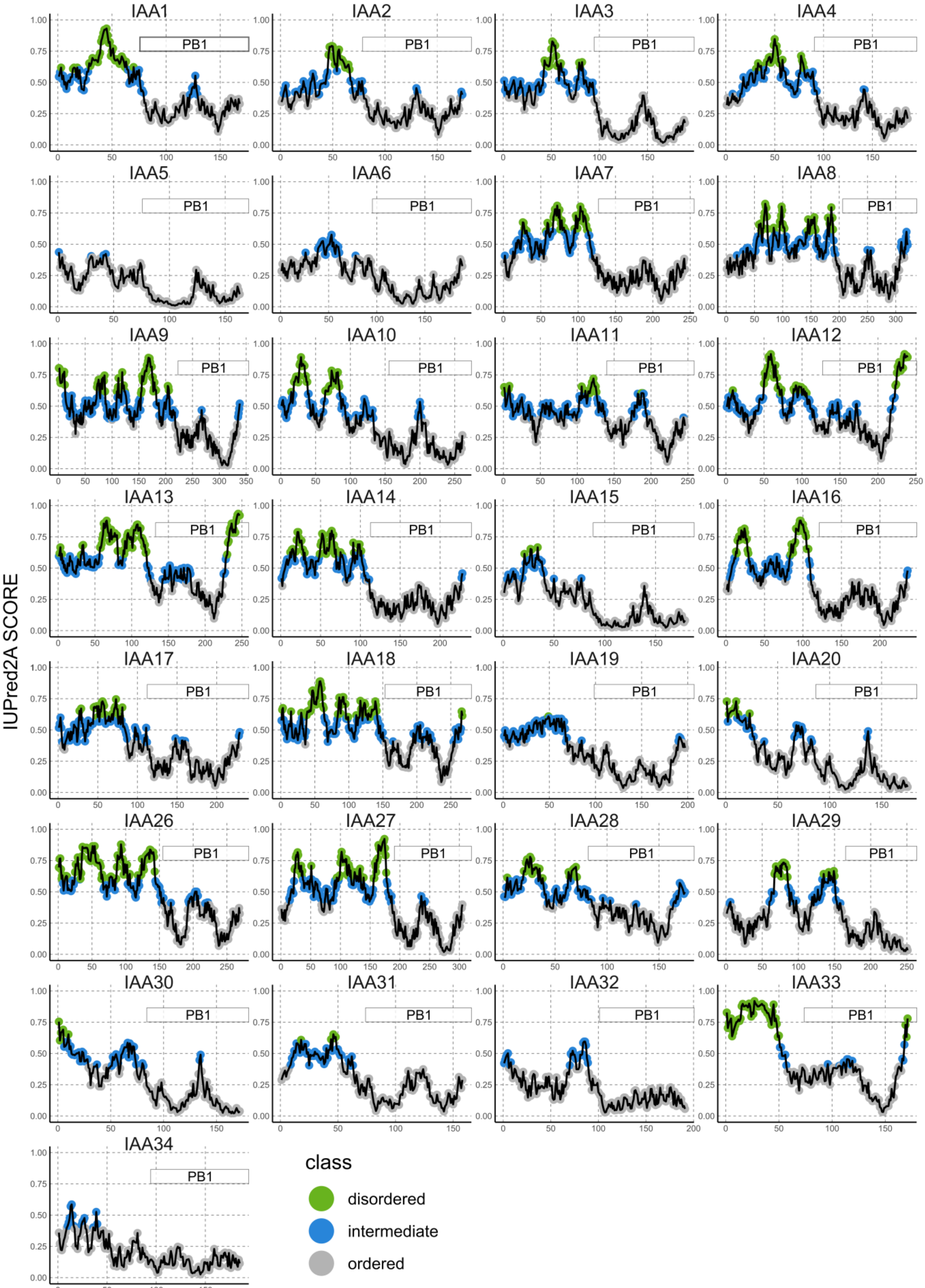


**Supplementary Figure 1| IUPred2A disorder prediction along the sequence of *Arabidopsis thaliana* AUX/IAA proteins.** The x-axis corresponds to the full length of each AUX/IAA protein sequence, and the y-axis shows the IUPred2A score for each amino acid (probability between 0-1). Amino acid residues are colored according to their disorder probability (disordered: ≥ 0.6, green; intermediate: 0.4-0.6, blue and ordered: ≤0.4, gray). The resolved, ordered PB1 domain is located along the sequence, as indicated, starting with the conserved VKV motif.

### Supp. Fig. 2

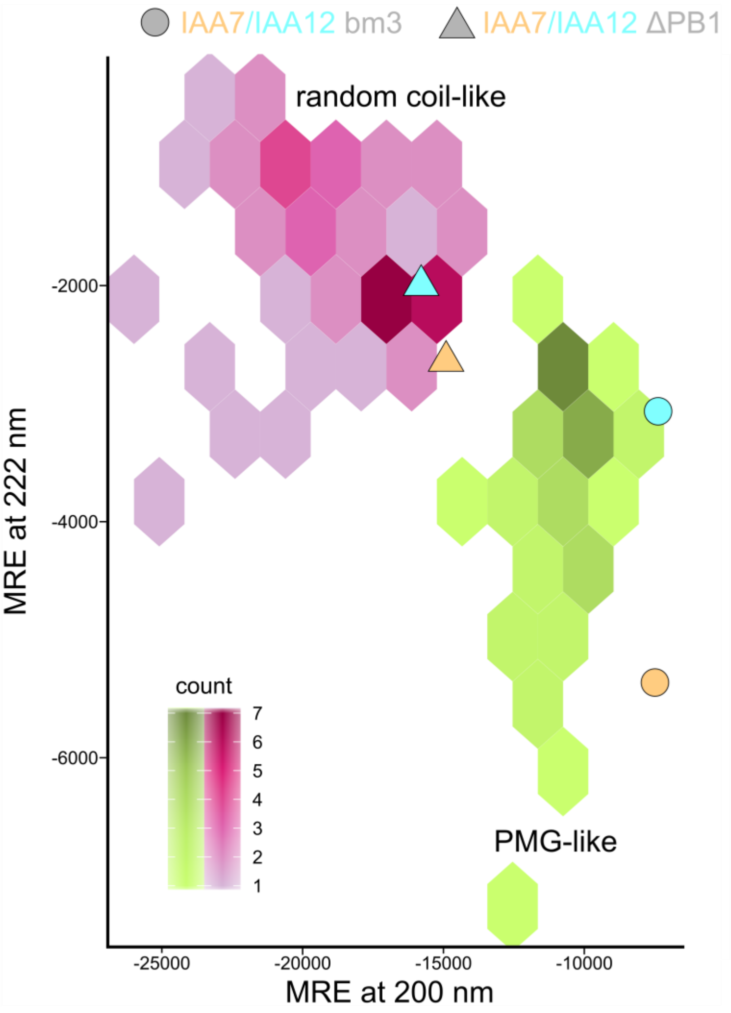


**Supplementary Figure 2| Classification of IAA7 and IAA12 variants according to their CD spectra.** CD spectral data classifies IAA7 (light orange) and IAA12 (aquamarine) as PMG-like proteins with random coil elements in their N-terminal half[1-3]. Molar residual ellipticity (MRE) at 200 nm and 222 nm is shown for the specified AUX/IAA protein variants on top of hexagonal binned reference proteins, with either unfolded, random coil-like proteins (purple) or premolten globule-like (PMG-like; green) proteins. Truncated versions (triangles, ΔPB1) lack the conserved folded PB1 domain. IAA7bm3 and IAA12bm3 variants (circles) carry 3 amino acid exchanges in their PB1 domain to render them oligomerization deficient.

### Supp. Fig. 3

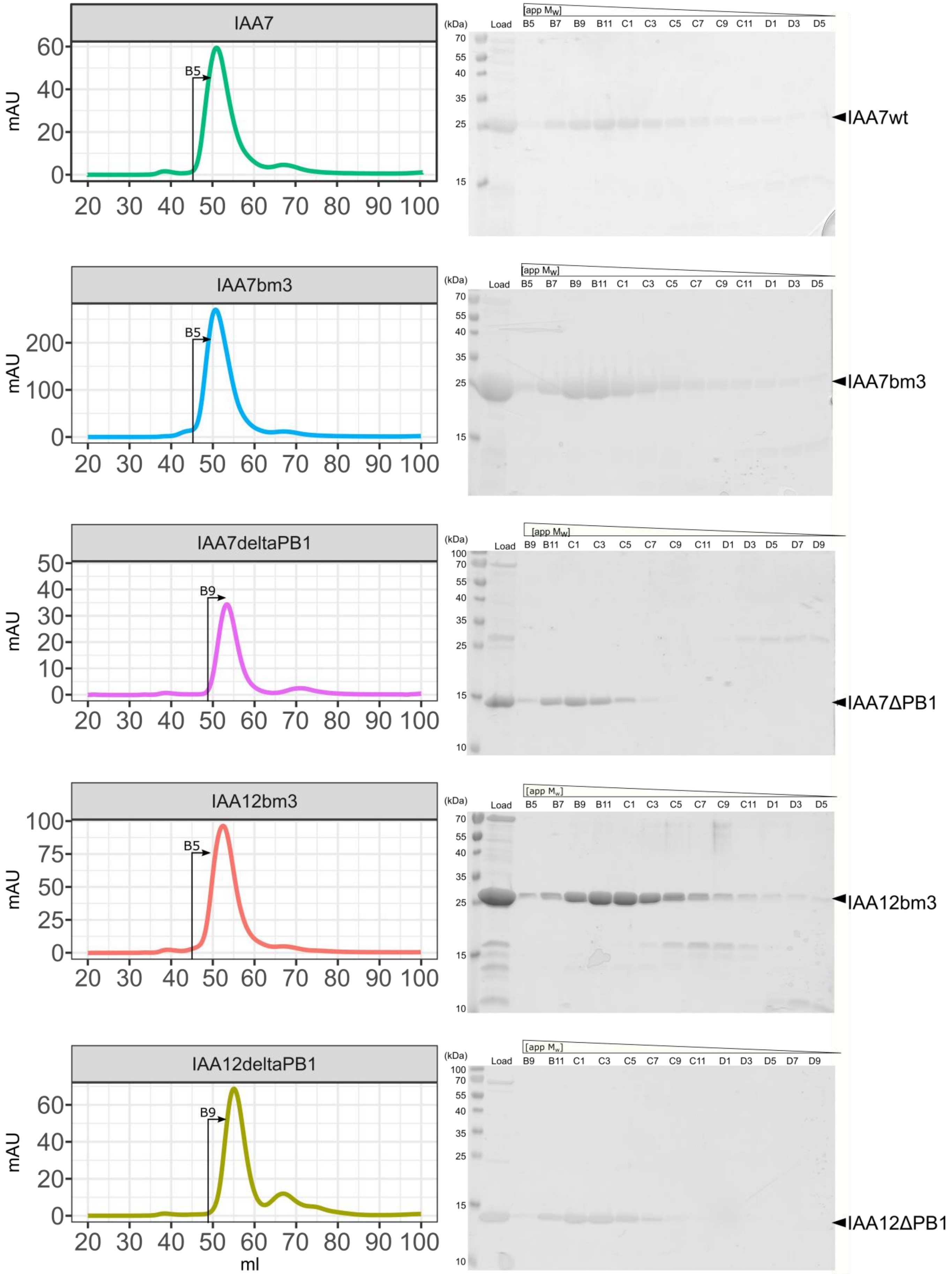


**Supplementary Figure 3| Representative size exclusion chromatography runs for the untagged AUX/IAA protein variants.** Elution profiles were obtained from semi-preparative size exclusion chromatography runs on a calibrated HiPrep 16/60 Sephacryl S100 High Resolution column (left panels). Indicated is the first fraction analyzed by SDS-PAGE shown in the right panels. Impurities could be separated from the protein of interest (indicated).

### Supp. Fig. 4

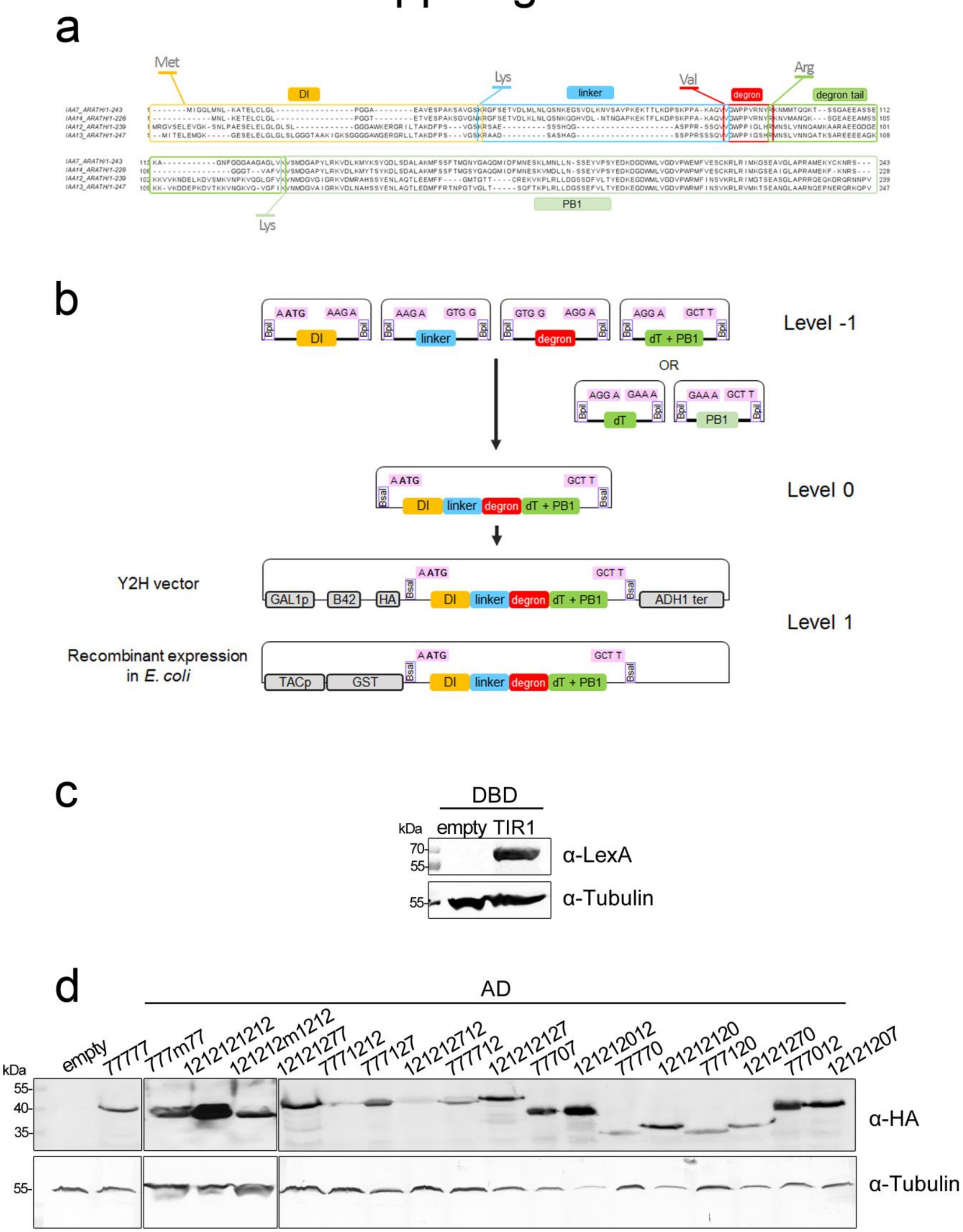


**Supplementary Figure 4| Design principle for 4- and 5-module AUX/IAA chimeras. a,** Sequence alignment of IAA7, IAA14, IAA12, IAA13 from *Arabidopsis thaliana* showing conserved amino acid residues selected as start and end of each module: DI (orange), linker (blue), core degron (red), degron tail (dark green) and the PB1 domain (light green). Conserved amino acids used as Golden Gate assembly sites are highlighted. **b,** Golden Gate cloning strategy to assemble level -1, 0, and 1 constructs for either yeast-two hybrid assays or recombinant *E.coli* expression as GST-fusion proteins using *Bpi*I and *Bsa*I restriction enzymes. **c-d,** Immunoblots for LexA-DBD-tagged TIR1 (**c**) and HA-tagged AUX/IAA chimeras (**d**) from haploid yeast cells grown in Gal/Raff –Trp or Gal/Raff –Ura –His medium, respectively. Detection was carried out using anti-LexA, anti-HA (F7), and anti-tubulin (loading control) antibodies.

### Supp. Fig. 5

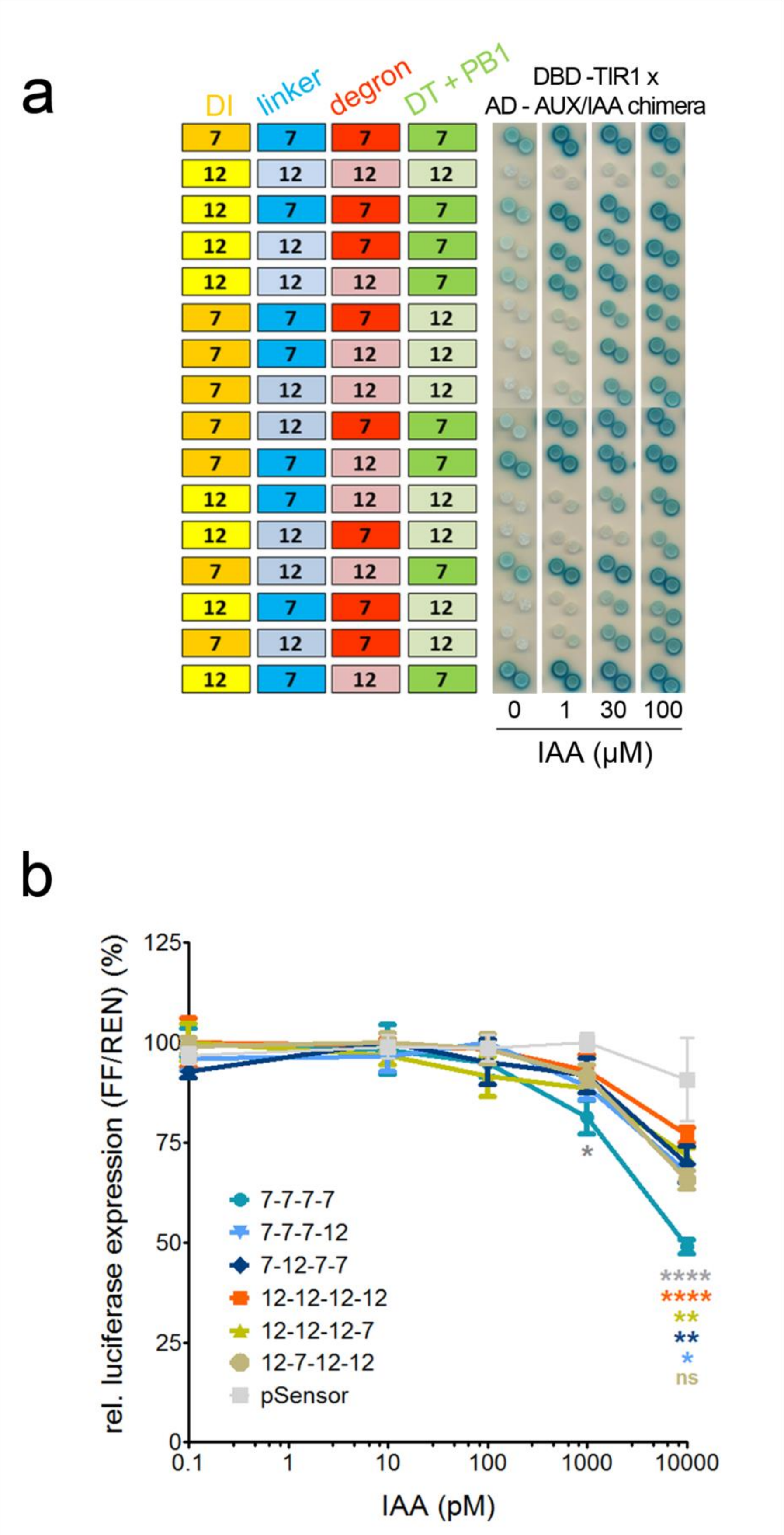


**Supplementary Figure 5| Design of 4-module chimeras, where module 4 consists of the degron tail and the PB1 domain of IAA7 or IAA12 combined. a,** Yeast two hybrid assay shows auxin-dependent interaction of TIR1 and chimeric AUX/IAAs is strongly driven by the presence of the IAA7 degron tail, and the PB1 domain -containing module. **b,** Ratiometric luminescent biosensor[4] to track degradation of 4-module AUX/IAA chimeric proteins in *Arabidopsis* protoplasts.

### Supp. Fig. 6

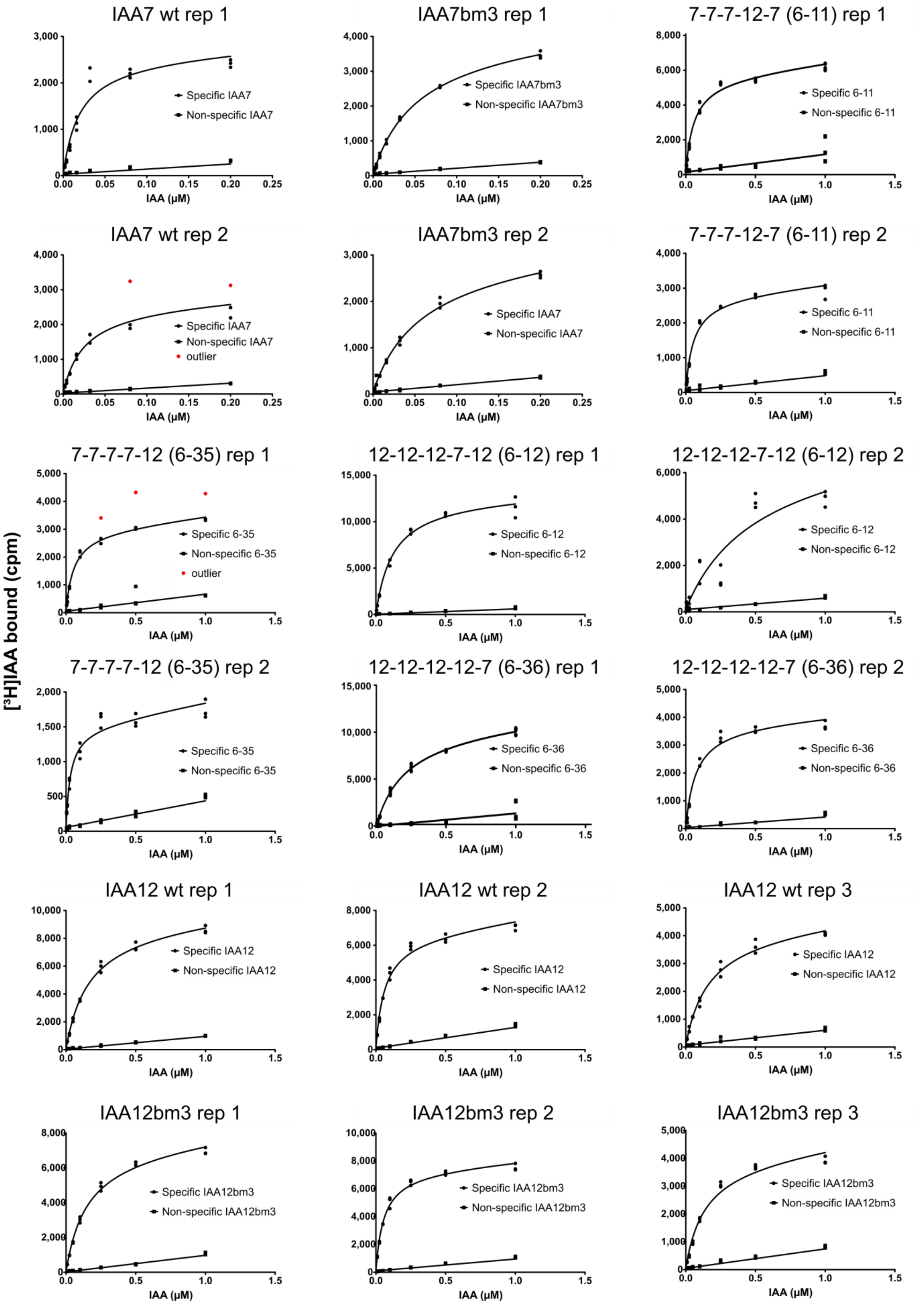


**Supplementary Figure 6| Single non-normalized [³H] IAA radioligand binding curves.** Single binding curves for each AUX/IAA variant and chimeric construct. Datapoints of each [³H] IAA concentration are shown as individual points for each technical replica (circles) together with non-specific binding in the presence of 2 mM cold IAA (squares). If present, outliers are marked in red.

Supp. Fig. 7

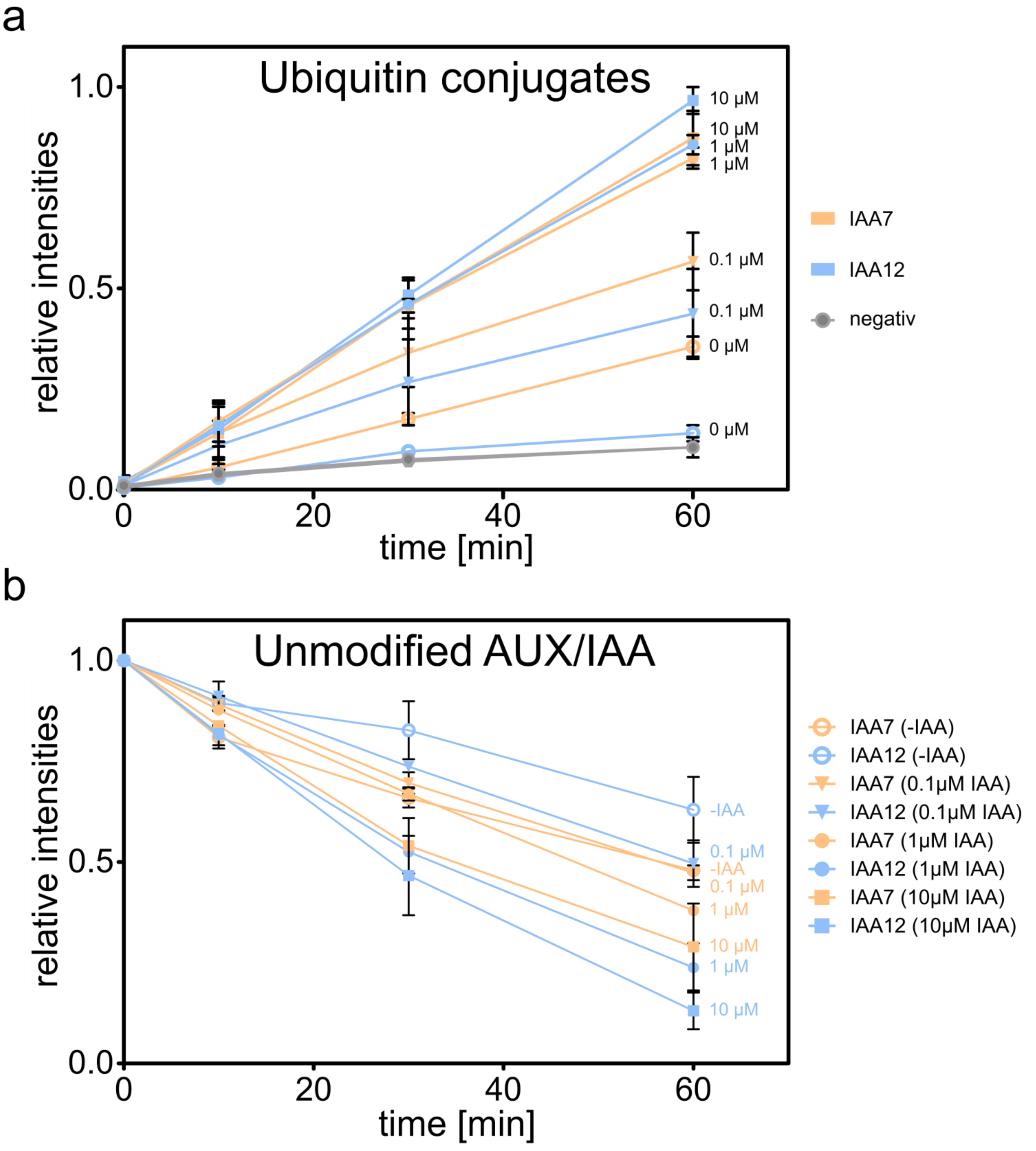


**Supplementary Figure 7| Quantification of auxin- and time-dependent ubiquitylation of IAA7 and IAA12. a,** Increase in ubiquitin conjugates over time measured as the in-gel ubiquitin-fluorescein signal intensity above the ubiquitin-modified Cullin1 (asterisk, **Figure 3a).** Signal was normalized by the strongest signal (IAA12, 10 µM IAA). **b,** Decrease of unmodified GST-AUX/IAA protein signal after immunoblotting detected by an Alexa Fluor Plus 647-coupled secondary antibody. Signals were normalized to the intensities at time point “0“. Depicted are mean values from three independent experiments with standard deviation as error bars. Results for GST-IAA7 and GST-IAA12 are depicted in light orange and light blue, respectively.

### Supp. Fig. 8

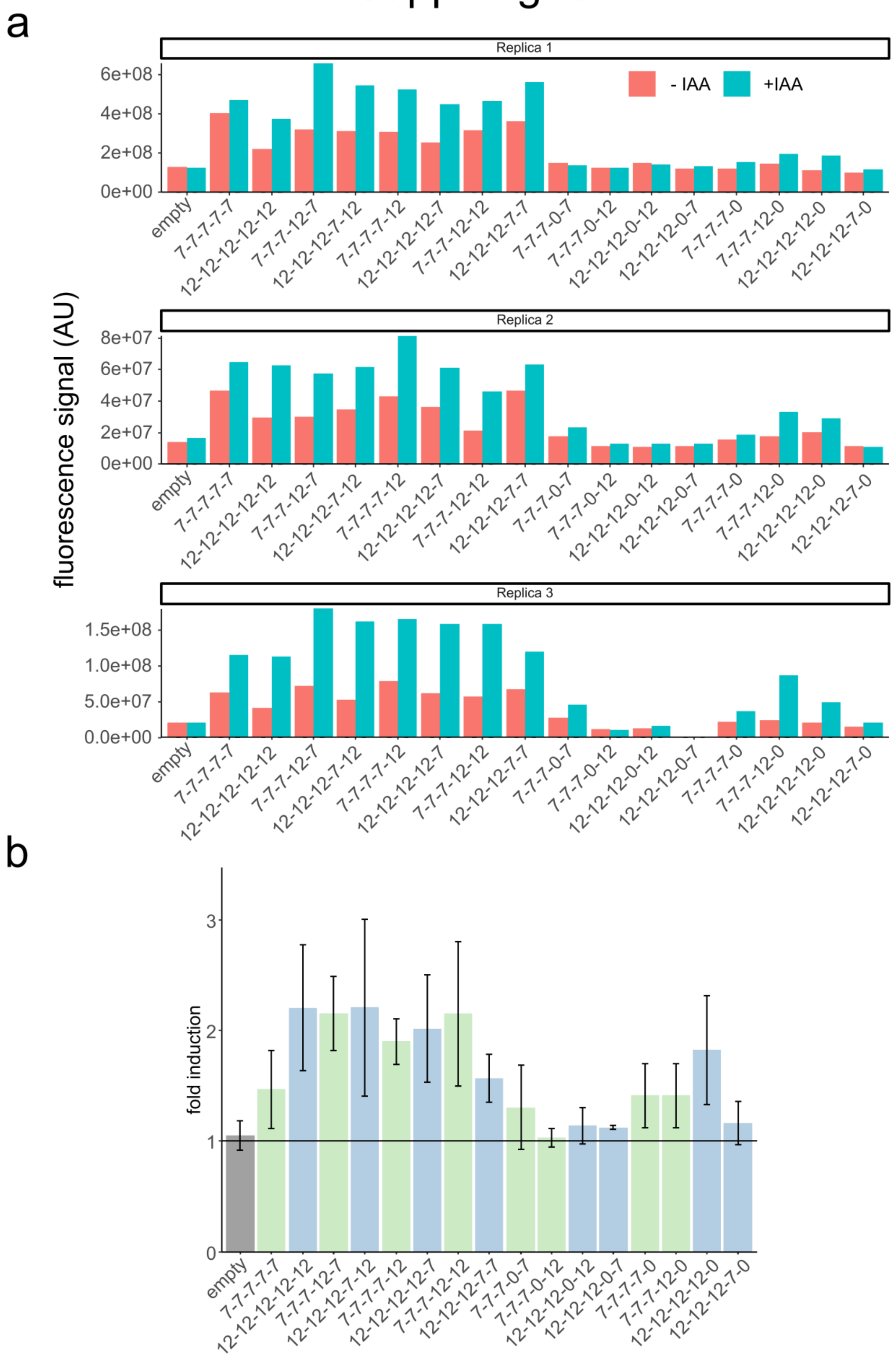


**Supplementary Figure 8| Quantification of auxin-triggered chimera ubiquitylation.** As in Supplemental Figure 6 ubiquitin conjugates on chimeric AUX/IAAs were measured via fluorescein signal intensities in the presence (teal) or absence (salmon) of auxin (IAA) after 1 h reaction time. **a,** Raw signal intensities for each individual replica. **b,** Auxin-triggered fold induction of chimera ubiquitylation as mean values with standard deviation using data from **a**. Chimeras consisting mainly of IAA7 (pale green) or IAA12 (light blue) modules are displayed.

Supp. Fig. 9

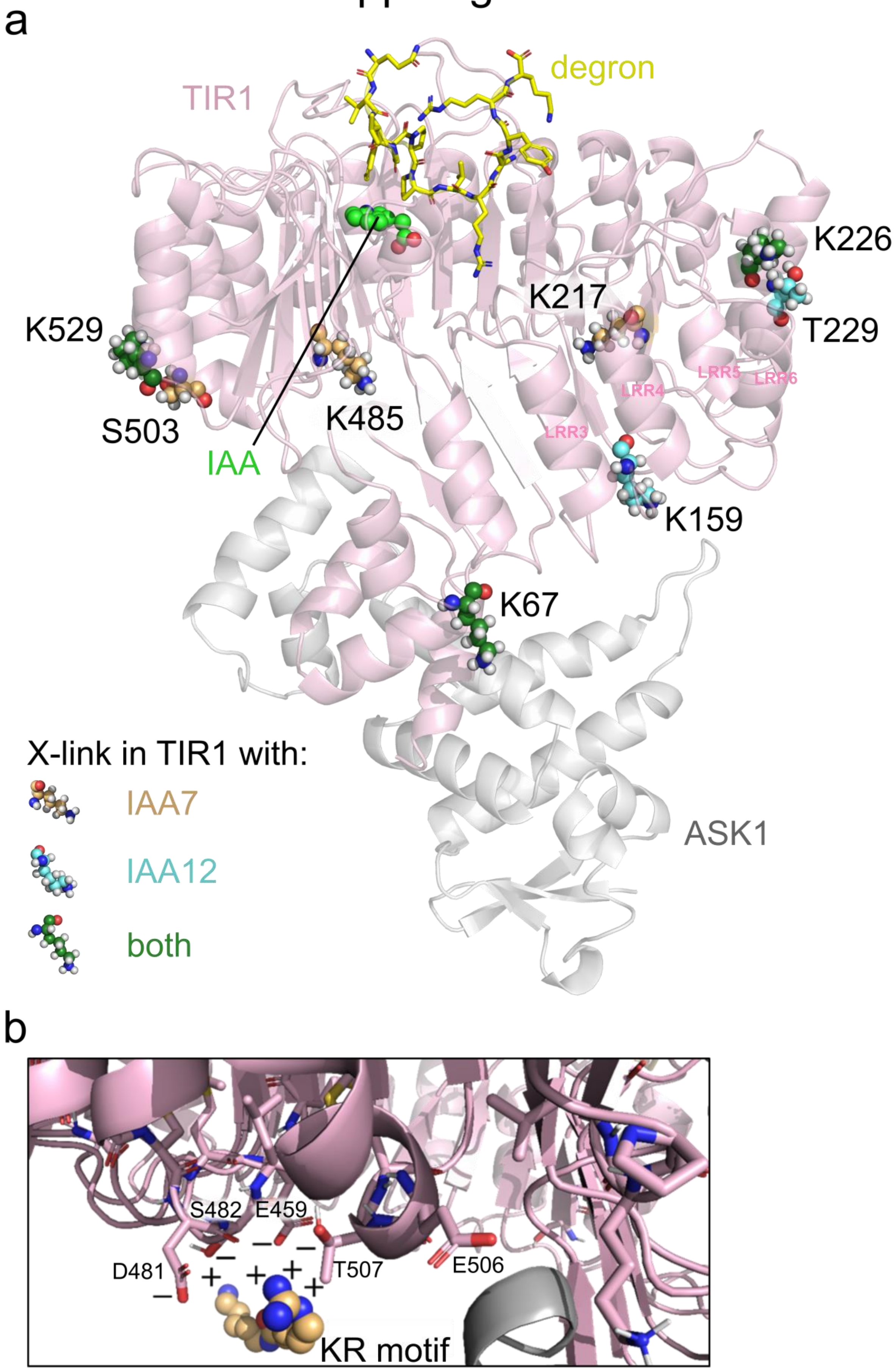


**Supplementary Figure 9| Crosslinked residues in TIR1 either with IAA7, IAA12 or both. a,** Depicted is the crystal structure of ASK1·TIR1·auxin·IAA7 degron (2P1Q, gray, light pink) with highlighted residues found to be crosslinked with either IAA7 (light orange), IAA12 (aquamarine) or both (green) shown as spheres. Leucine-rich repeats carrying PB1 domain-interacting are labeled. **b,** Patch enriched with negative charge potential, close to KR motif-cross-linked residues, acting as a plausible interaction site.

### Supp. Fig. 10

a

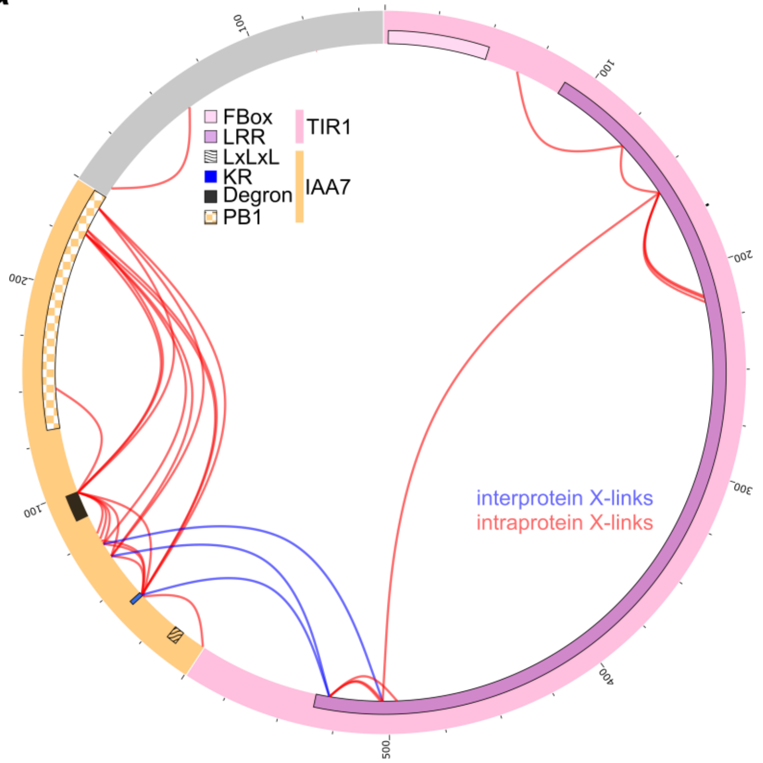


b

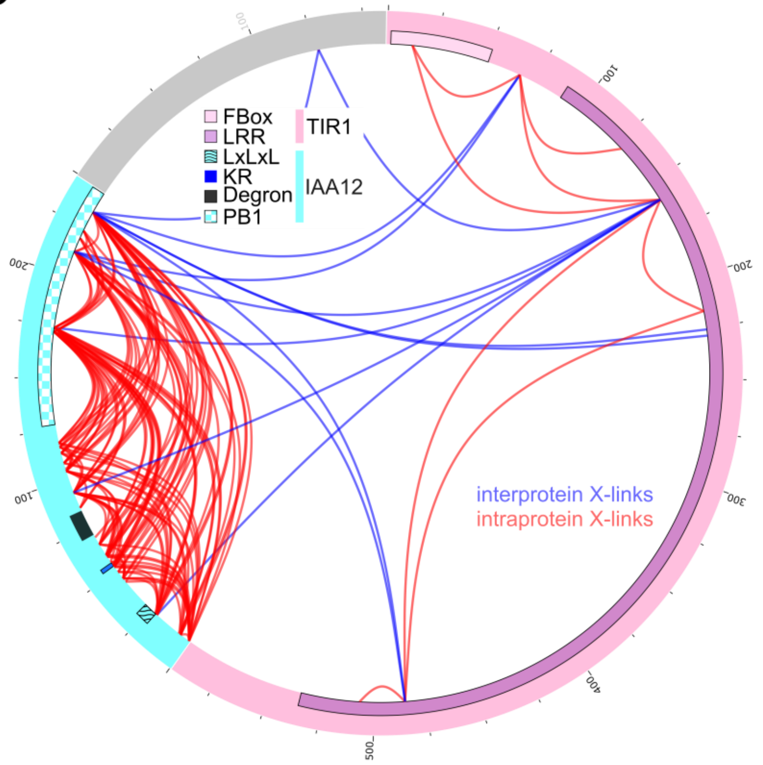


**Supplementary Figure 10| Crosslinks identified in ASK1·TIR1 and AUX/IAAs in the absence of auxin.** Displayed are all crosslinks within (intra-protein, red) or in between (inter-protein, blue) ASK1 (gray), TIR1 (light pink) and IAA7 (light orange, **a**) or IAA12 (aquamarine, **b**) as connecting lines along the circular depicted amino acid sequence. Lines correspond to all crosslinked peptides collected from multiple replica.

Supp. Fig. 11

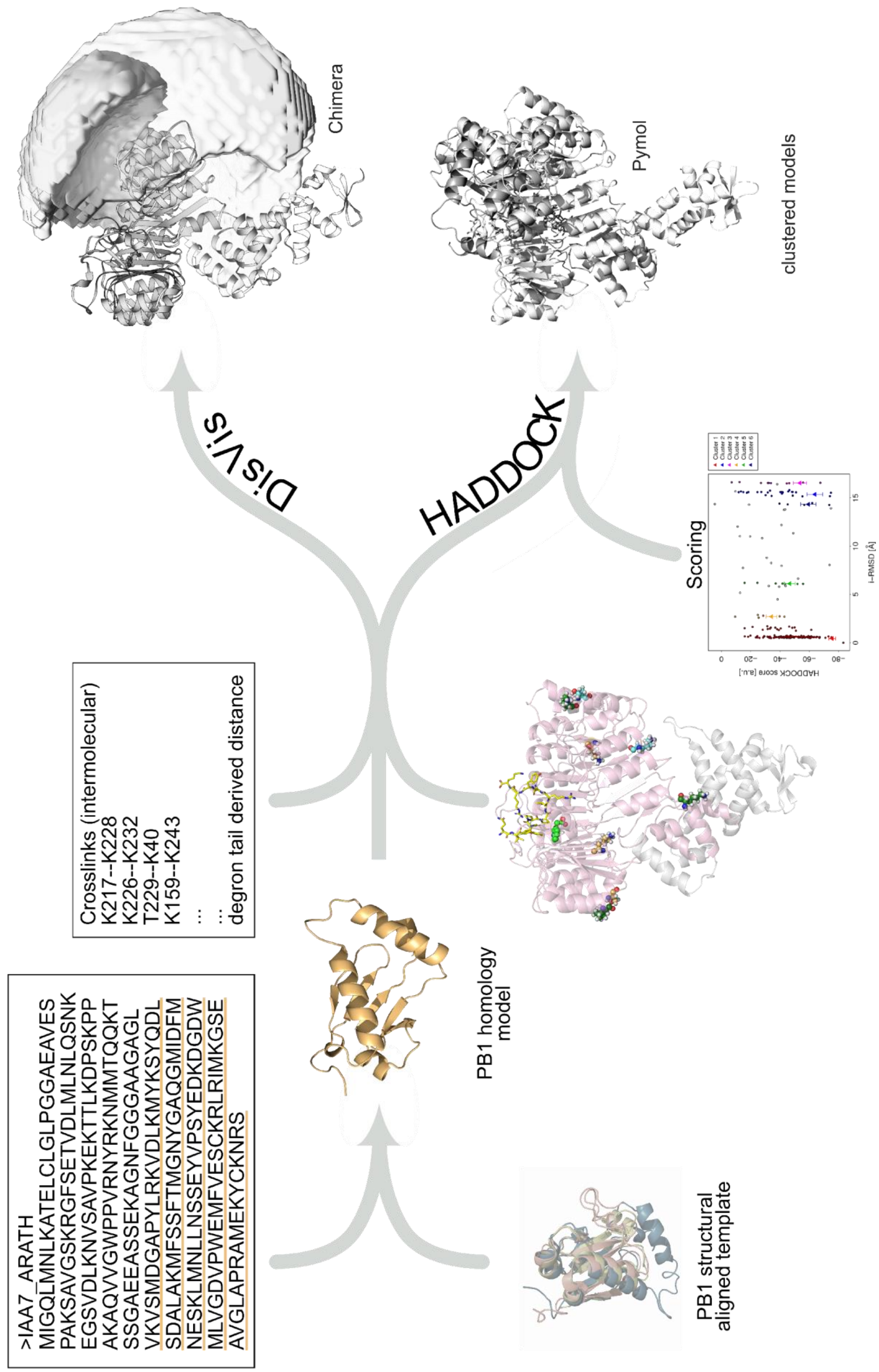


**Supplementary Figure 11| Workflow for cross-linking-based docking using HADDOCK.** Homology models from *At*IAA7 and IAA12 PB1 domains were created using multi-template-based comparative modelling with MODELLER. Docking models using HADDOCK were generated by docking the PB1 homology models on the modified ASK1·TIR1·auxin·degron crystal structure (2P1Q) using as distant restraints the cross-linking information and the degron tail length. Potential conformational space was visualized via DisVis.

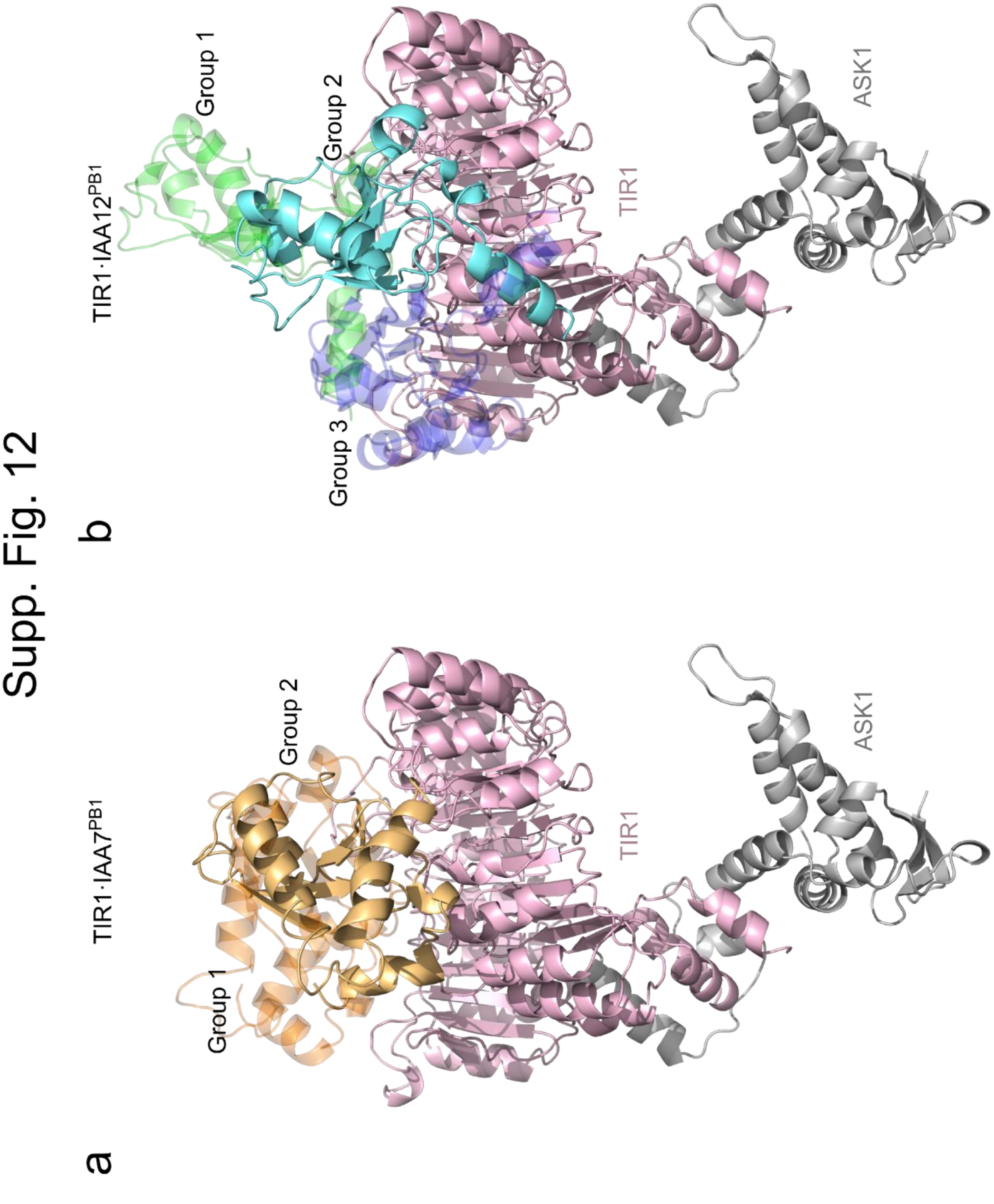


**Supplementary Figure 12| HADDOCK docking using crosslinking data restrictions generated structural models of TIR1·AUX/IAA PB1 complexes.** A representative structure for each group of TIR1·IAA7 PB1 (**a**) (group 1 (dark orange), group 2 (light orange)); and TIR1·IAA12 PB1 (**b**) (group 1 (green), group 2 (aquamarine), group 3 (dark blue)) HADDOCK models are shown.

### Supp. Fig.13

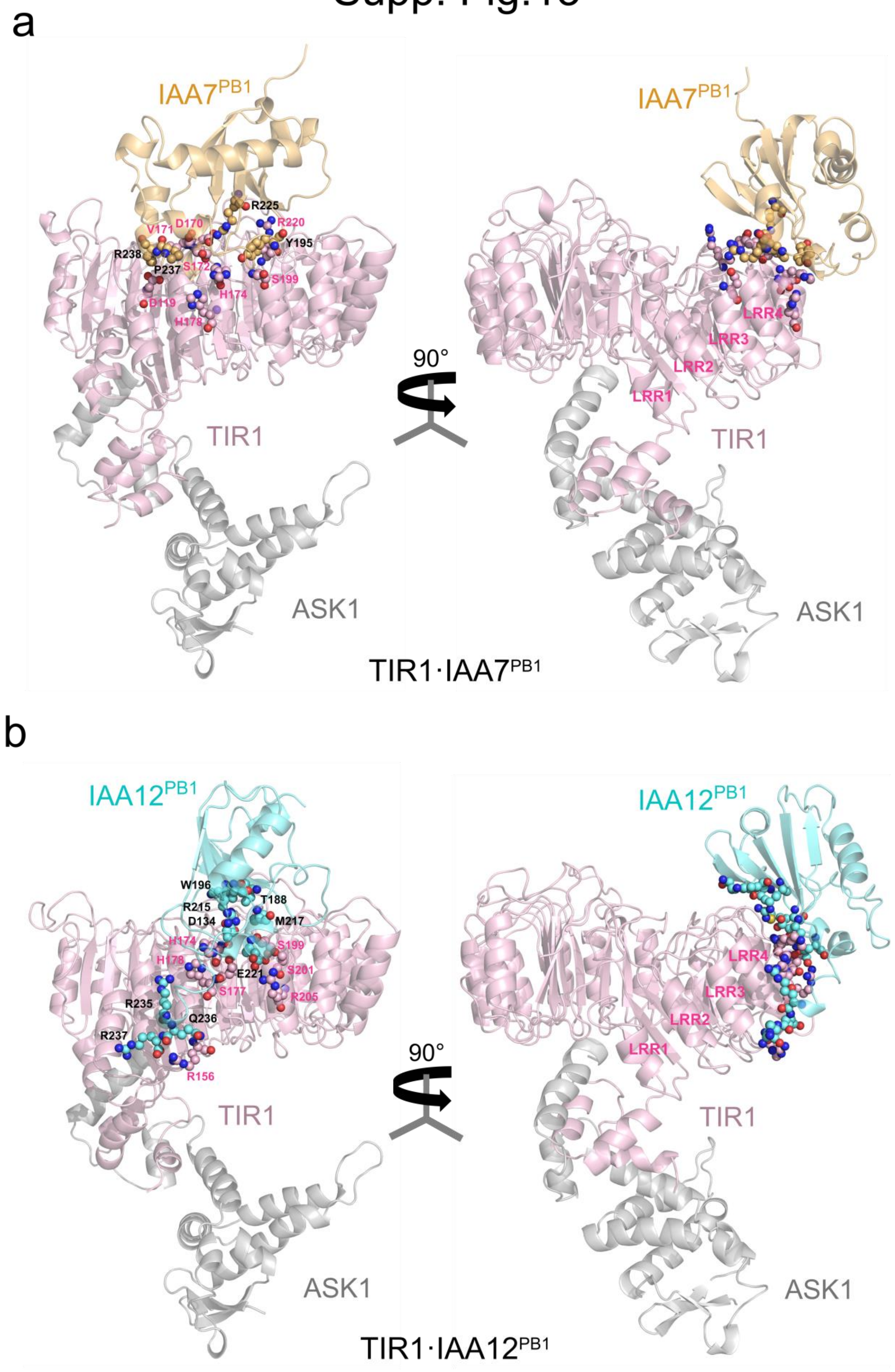


**Supplementary Figure 13| Molecular dynamics (MD) simulations revealed the most energetically favorable model from HADDOCK-based docking. a-b,** PB1 domains from both IAA7 and IAA12 are positioned over TIR1 interacting with residues from leucine-rich-repeat 3-6 (LRR3-6). Energetically relevant residues (small spheres) from TIR1 (light pink), IAA7 PB1 (light orange), and IAA12 PB1 (aquamarine) domains for complex stabilization are located in the TIR1·AUX/IAA PB1 interface.

### Supp. Fig. 14

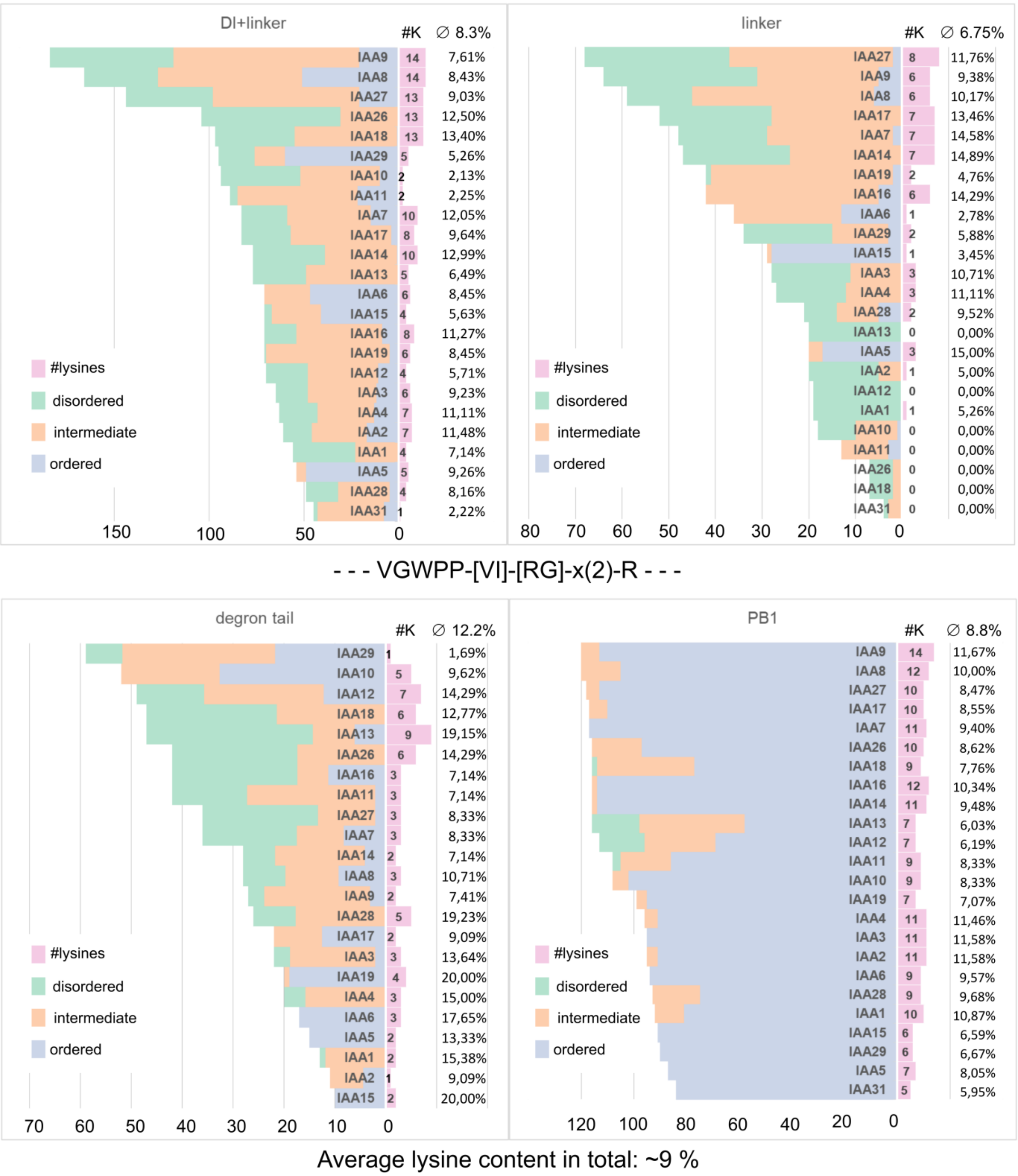


**Supplementary Figure 14| Disorder probability and lysine content in different regions of the canonical *At*AUX/IAA proteins.** IUPred2A-based prediction for disordered (green), intermediate (orange) and ordered (blue) amino acid residues is shown. Length in AUX/IAA IDRs partially correlate with lysine content and/or disorder. AUX/IAAs with less than 5 lysine residues (pink) in the degron tail show increased lysine content in the PB1 domain (≥10). AUX/IAA degron tails are enriched in ubiquitin acceptors sites (lysine residues, average 12.2% of total residues).

**Suppl. Table 2. Crosslinking-based docking by HADDOCK**

| Input | # grouped refined structures | group | # structures per group | HADDOCK scores | Buried surface | Van der Waals energy | Electrostatic energy | Restraint violation |
|---|---|---|---|---|---|---|---|---|
| ASK1·TIR1·IAA7 without degron tail restraint | 175 | 1 | 124 | -75.7 +/- 4.6 | 1390.6 +/- 29.4 | -37.9 +/- 3.1 | -279.1 +/- 15.2 | 2.5 +/- 0.72 |
| | | 2 | 24 | -63.8 +/- 10.8 | 1503.0 +/- 127.2 | -47.9 +/- 8.8 | -214.5 +/- 95.6 | 2.7 +/- 1.50 |
| | | 6 | 5 | -59.4 +/- 10.2 | 1627.6 +/- 95.8 | -46.7 +/- 6.5 | -210.5 +/- 15.6 | 2.3 +/- 1.02 |
| | | 3 | 10 | -53.6 +/- 9.1 | 1451.2 +/- 59.6 | -32.4 +/- 6.2 | -265.6 +/- 60.2 | 1.6 +/- 0.53 |
| | | 5 | 6 | -46.7 +/- 7.6 | 1019.5 +/- 31.2 | -38.8 +/- 3.9 | -127.3 +/- 32.5 | 2.5 +/- 0.56 |
| | | 4 | 6 | -34.2 +/- 7.4 | 1031.6 +/- 41.0 | -22.7 +/- 2.5 | -234.4 +/- 27.5 | 2.1 +/- 0.54 |
| ASK1·TIR1·IAA7 with degron tail restraint | 193 | 2 | 72 | -89.1 +/- 2.5 | 1689.6 +/- 146.4 | -43.5 +/- 3.0 | -419.7 +/- 33.1 | 3.2 +/- 0.84 |
| | | 1 | 121 | -66.7 +/- 9.3 | 1395.2 +/- 162.7 | -33.5 +/- 3.8 | -272.7 +/- 39.0 | 1.6 +/- 0.36 |
| ASK1·TIR1·IAA12 without degron tail restraint | 132 in 13 groups | 2 | 18 | -76.3 +/- 10.0 | 1588.8 +/- 70.0 | -38.1 +/- 4.8 | -293.4 +/- 45.0 | 2.5 +/- 0.44 |
| | | 1 | 47 | -67.1 +/- 12.1 | 990.6 +/- 15.7 | -30.3 +/- 1.6 | -260.0 +/- 31.6 | 1.9 +/- 0.62 |
| | | 4 | 9 | -66.8 +/- 4.0 | 1548.8 +/- 140.0 | -44.8 +/- 11.7 | -257.4 +/- 46.3 | 2.6 +/- 0.39 |
| | | 9 | 5 | -66.5 +/- 10.6 | 1090.0 +/- 38.5 | -23.9 +/- 1.4 | -381.9 +/- 33.2 | 3.1 +/- 1.26 |
| | | 8 | 5 | -61.5 +/- 22.4 | 1218.9 +/- 164.4 | -27.5 +/- 5.4 | -404.8 +/- 62.1 | 2.5 +/- 0.69 |
| | | 11 | 5 | -51.9 +/- 4.1 | 1425.8 +/- 80.1 | -29.5 +/- 2.0 | -357.4 +/- 35.3 | 2.0 +/- 0.69 |
| | | 10 | 5 | -45.7 +/- 5.6 | 1434.8 +/- 70.1 | -38.3 +/- 3.4 | -230.4 +/- 29.4 | 1.9 +/- 0.33 |
| | | 7 | 6 | -41.5 +/- 3.5 | 1129.3 +/- 38.6 | -27.1 +/- 2.5 | -267.7 +/- 10.3 | 2.0 +/- 0.99 |
| | | 3 | 10 | -35.6 +/- 3.7 | 1193.6 +/- 31.5 | -30.6 +/- 1.7 | -89.6 +/- 21.8 | 2.9 +/- 0.74 |
| | | 13 | 4 | -33.1 +/- 8.6 | 1161.6 +/- 120.1 | -24.3 +/- 4.5 | -207.1 +/- 21.8 | 1.9 +/- 0.42 |
| ASK1·TIR1·IAA12 with degron tail restraint | 196 | 1 | 187 | -94.2 +/- 9.9 | 1619.0 +/- 85.6 | -35.4 +/- 2.4 | -469.4 +/- 48.7 | 1.8 +/- 0.30 |
| | | 3 | 4 | -59.8 +/- 14.5 | 1541.9 +/- 116.0 | -40.6 +/- 2.8 | -252.6 +/- 58.2 | 2.8 +/- 0.44 |
| | | 2 | 5 | -53.9 +/- 10.2 | 1429.8 +/- 104.0 | -38.0 +/- 4.5 | -229.1 +/- 35.9 | 3.0 +/- 1.58 |

#### Supp. Table 3. Per-residue energy contributions to the formation of the TIR1·PB1 complexes

| Complexes | TIR1 Residues | Conservation within 21 TIR1/AFB-like proteins | prEFED protocol | | CAS protocol | |
|---|---|---|---|---|---|---|
| | | | $\Delta G_{sc}$ (kcal/mol) $GB^{OBC1}$ | $\Delta G_{sc}$ (kcal/mol) $GB^{OBC2}$ | ΔΔG (kcal/mol) $GB^{OBC1}$ | ΔΔG (kcal/mol) $GB^{OBC2}$ |
| TIR1·IAA7 | D170 | 5 (+8) | -3.746 +/- 1.11 | -5.317 +/- 1.32 | -13.678 +/- 2.06 | -16.533 +/- 2.36 |
| | R220 | 9 (+4) | -6.543 +/- 1.35 | -6.975 +/- 1.46 | -9.585 +/- 2.29 | -10.623 +/- 2.54 |
| | D119 | 19 | -2.264 +/- 0.76 | -3.364 +/- 0.98 | -8.632 +/- 1.48 | -10.414 +/- 1.86 |
| | H174 | 2 | -4.902 +/- 0.85 | -5.418 +/- 0.97 | -7.387 +/- 1.57 | -8.255 +/- 1.78 |
| | S172 | 5 | -4.011 +/- 0.62 | -4.241 +/- 0.65 | -6.671 +/- 1.23 | -7.447 +/- 1.28 |
| | S199 | 2 | -3.103 +/- 1.23 | -3.375 +/- 1.30 | -4.558 +/- 2.04 | -5.032 +/- 2.18 |
| | H178 | 1 (+3) | -2.841 +/- 0.59 | -3.041 +/- 0.64 | -2.899 +/- 0.95 | -3.160 +/- 1.02 |
| | V171 | 1 | -2.536 +/- 0.49 | -2.278 +/- 0.51 | -2.483 +/- 0.90 | -2.099 +/- 0.95 |
| | E197 | 11 (+4) | 2.437 +/- 1.07 | 2.331 +/- 1.22 | -0.149 +/- 2.73 | -0.807 +/- 3.14 |
| | D146 | 8 (+2) | 1.988 +/- 0.89 | 1.959 +/- 0.88 | 0.709 +/- 1.34 | 0.385 +/- 1.55 |
| | K226 | 2 | 0.685 +/- 0.56 | 1.077 +/- 0.91 | - | - |
| TIR1·IAA12 | R205 | 6 (+2) | -7.99 +/- 0.90 | -7.662 +/- 1.00 | -15.253 +/- 1.84 | -16.056 +/- 2.03 |
| | R156 | 19 (+2) | -7.44 +/- 0.97 | -7.555 +/- 1.05 | -11.491 +/- 1.88 | -12.437 +/- 1.99 |
| | H174 | 2 | -4.267 +/- 1.52 | -5.195 +/- 1.98 | -9.27 +/- 2.46 | -10.690 +/- 2.89 |
| | S201 | 2 | -4.058 +/- 0.60 | -4.508 +/- 0.64 | -7.689 +/- 1.15 | -8.963 +/- 1.23 |
| | S199 | 2 | -3.942 +/- 1.15 | -4.439 +/- 1.18 | -7.448 +/- 2.19 | -8.731 +/- 2.36 |
| | H178 | 1 (+3) | -1.706 +/- 0.93 | -2.27 +/- 1.15 | -4.349 +/- 1.78 | -5.392 +/- 2.09 |
| | S177 | 11 (+2) | -1.521 +/- 1.32 | -1.738 +/- 1.40 | -2.833 +/- 2.32 | -3.307 +/- 2.54 |
| | K130 | 2 (+1) | -1.041 +/- 1.27 | -0.877 +/- 1.18 | -1.745 +/- 2.72 | -1.658 +/- 2.82 |
| | S196 | 5 (+2) | -1.247 +/- 0.41 | -1.281 +/- 0.47 | -1.286 +/-0.66 | -1.423 +/- 0.78 |
| | V171 | 1 | -1.784 +/- 0.38 | -1.548 +/- 0.41 | -0.767 +/- 0.87 | -0.459 +/- 0.94 |
| | A153 | 5 | -1.259 +/- 0.24 | -1.286 +/- 0.24 | - | - |
| | D170 | 5 (+8) | 0.459 +/- 0.16 | 0.059 +/- 0.23 | - | - |

**Supplementary Table 3| Energy contribution of single amino acids to TIR1·AUX/IAA complex formation.** Conservation of residues was checked in TIR1/AFB-like proteins in *Arabidopsis thaliana* (uniprot ID: Q570C0, Q9ZR12, Q9LW29, Q9LPW7, A0A178UVM5, A0A178UB83), *Selaginella moellendorffii* (uniprot ID: D8RF91, D8SDE6, D8SG63, D8R5Z3), *Physcomitrella patens* (uniprot ID: A9SYG2, A9TAY1, A9T980, A9SZ50, A9TE08, A9TP16), *Oryza sativa* (uniprot ID: Q0DKP3, Q7XVM8, Q2R3K5, Q8H7P5) and *Marchantia polymorpha* (uniprot ID: A0A2R6WBN4).
